# Asymmetric flagellar filaments shape *Borreliella burgdorferi* morphology and motility

**DOI:** 10.64898/2026.10.15.759423

**Authors:** Jiaqi Wang, Kai Zhang, Wangbiao Guo, Syed Z. Sultan, Aaron Ye, Anthony Low, Md A. Motaleb, Chunhao Li, Jun Liu

**Author notes:** These authors contributed equally to this work (Jiaqi Wang, Kai Zhang, Wangbiao Guo).

## Abstract

The Lyme disease spirochete *Borreliella burgdorferi* possesses bipolar periplasmic flagella that are essential for its distinctive flat-wave morphology, motility, and infectivity. However, how the molecular architecture of these flagella contributes to spirochetal morphology and motility remains poorly understood. Here, using *in situ* single-particle cryo-electron microscopy, we determine near-atomic structures of native periplasmic flagellar filaments and reveal two distinct architectures: half-sheathed and unsheathed. We show that the spirochete-specific protein FlaA forms an asymmetric sheath along the convex surface of the conserved FlaB filament core, with sheath assembly restricted to the proximal filament region near the flagellar motor. Atomic models reveal that FlaA anchors to the FlaB core through specific interactions with two surface-exposed loops, while genetic and motility analyses demonstrate distinct roles for these loops in maintaining cell morphology and motility. Structural comparisons reveal that the FlaB core possesses intrinsic curvature and supercoiling, which are substantially amplified by asymmetric FlaA assembly. These findings support a model in which localized FlaA sheath assembly reinforces the proximal filament and potentially modulates its mechanical properties. Together, our results uncover a previously unrecognized mechanism by which asymmetric sheath assembly enhances flagellar supercoiling, providing a molecular basis for understanding the distinctive morphology and motility of spirochetes.

**Significance:** The Lyme disease spirochete *Borreliella burgdorferi* relies on periplasmic flagella for its distinctive morphology and motility, but how filament architecture contributes to these properties remains unclear. Using *in situ* cryo-electron microscopy, we reveal that the spirochete-specific protein FlaA forms an asymmetric sheath around the FlaB filament core exclusively near the cell poles. FlaA anchors to flexible surface loops of FlaB through specific molecular interactions, while genetic analyses reveal distinct roles for these loops in maintaining cell morphology and motility. Asymmetric FlaA assembly amplifies the intrinsic supercoiling of the FlaB core, potentially modifying filament mechanics near the motor. These findings reveal how localized, asymmetric sheath assembly contributes to the distinctive morphology and motility of spirochetes.

## Main

*Borreliella burgdorferi*, the causative agent of Lyme disease, has evolved a unique flat-wave morphology and motility that enable the spirochete to navigate through complex host tissues and establish infection (1–3). Unlike externally flagellated bacteria, *B. burgdorferi* propels itself using periplasmic flagella, which are bundles of flagellar filaments located within the periplasmic space and wrapped around the cell cylinder between the inner and outer membranes (4–6). Rotation of these periplasmic flagella generates propagating waves along the cell body, producing a corkscrew-like mode of locomotion that is highly effective in viscous environments such as connective tissues and interstitial fluid (2, 6–8). Disruption of flagellar assembly or rotation renders *B. burgdorferi* non-motile, unable to migrate out of ticks to infect mice, and non-infectious in murine models, demonstrating that periplasmic flagella are not merely crucial for motility but are essential virulence determinants (9–11). Beyond driving motility, periplasmic flagella also play a skeletal role in maintaining the characteristic flat-wave morphology of *B. burgdorferi*, a feature unique among flagellated bacteria (6, 12, 13).

Each periplasmic flagellum is anchored near one cell pole through a membrane-embedded motor and extends toward the opposite pole, with the filament wrapping around the cell body. The *B. burgdorferi* flagellar filament is a heteropolymeric assembly composed of two proteins: FlaA and FlaB (14, 15). FlaB belongs to the highly conserved bacterial flagellin superfamily but is considerably smaller than enteric flagellins like *Salmonella* FliC (16) because it lacks the surface-exposed D2 and D3 domains that project from canonical flagellar filaments (17, 18). This architecture, recently confirmed at near-atomic resolution in *Treponema denticola*, produces a filament core approximately 12.5 nm in diameter that is well suited to the confined periplasmic space (19). In contrast to canonical bacterial flagellar systems, FlaA is a spirochete-specific flagellar protein that assembles on the FlaB outer surface to form the filament sheath (20, 21). *B. burgdorferi* encodes a single *flaB* and a single *flaA*, whereas *Treponema* and *Leptospira* species synthesize multiple FlaB paralogs together with distinct sheath proteins, including FcpA and FcpB (14, 22–24). Genetic studies have shown that deletion of *B. burgdorferi flaB* alters cell morphology and abolishes filament assembly and motility (6, 9). Although *flaA* mutants have not been fully characterized in *B. burgdorferi*, mutants lacking FlaA in other spirochetes, such as *Treponema* and *Brachyspira*, assemble unsheathed filaments and exhibit reduced motility, indicating that the sheath is dispensable for core filament assembly but essential for efficient motility in many spirochetes (19, 25, 26). Despite the well-established biological importance of periplasmic flagella, structural information of the native *B. burgdorferi* filament has remained elusive. Consequently, the molecular mechanisms governing FlaA–FlaB assembly and sheath organization and their contributions to filament architecture have remained unsolved in *B. burgdorferi*.

Here, we report near-atomic structures of the *B. burgdorferi* periplasmic flagellar filament determined by *in situ* single-particle cryo-electron microscopy (cryo-EM). These structures reveal the molecular architecture of the FlaB filament core, define the organization and interactions of the FlaA sheath with the FlaB core, and uncover a previously unrecognized asymmetric sheath architecture. By integrating near-atomic structure determination with genetic and motility analyses, we propose a model in which asymmetric sheath assembly modulates filament curvature and supercoiling near cell poles, establishing a molecular basis for understanding how the FlaA sheath regulates flagellar filament architecture and contributes to the distinctive cell morphology and motility of spirochetes.

## Results

### Characterization of the *B. burgdorferi* periplasmic flagellar filaments

The *B. burgdorferi* periplasmic flagellar filament is assembled by two proteins, FlaB (BB_0147) and FlaA (BB_0668) (1). FlaB lacks a signal peptide and is exported through the flagellar type III secretion system (fT3SS) (1), consistent with its position in the filament core, whereas FlaA carries a predicted signal peptide and reaches the periplasm through the Sec pathway (1), consistent with its position in the sheath (*SI Appendix*, Fig. S1*A*). Structure predictions show that FlaB adopts an α-helix-dominant structure with conserved D0 and D1 domains, while FlaA is composed of a β-rich domain with extended termini predicted with low confidence (*SI Appendix*, Fig. S1*B*).

To better understand the composition and organization of the *B. burgdorferi* periplasmic flagellar filament, we first examined the localization of sheath protein FlaA by immunofluorescence microscopy. FlaA labeling was confined to the cell poles, corresponding to the proximal regions of the periplasmic flagella adjacent to the flagellar motors (13, 27), whereas no detectable signal was observed along the remainder of the filament (Fig. 1*A–C*). These results indicate that FlaA is not uniformly distributed along the flagellar filament but is instead selectively incorporated into the pole-proximal region.

**Fig. 1.**
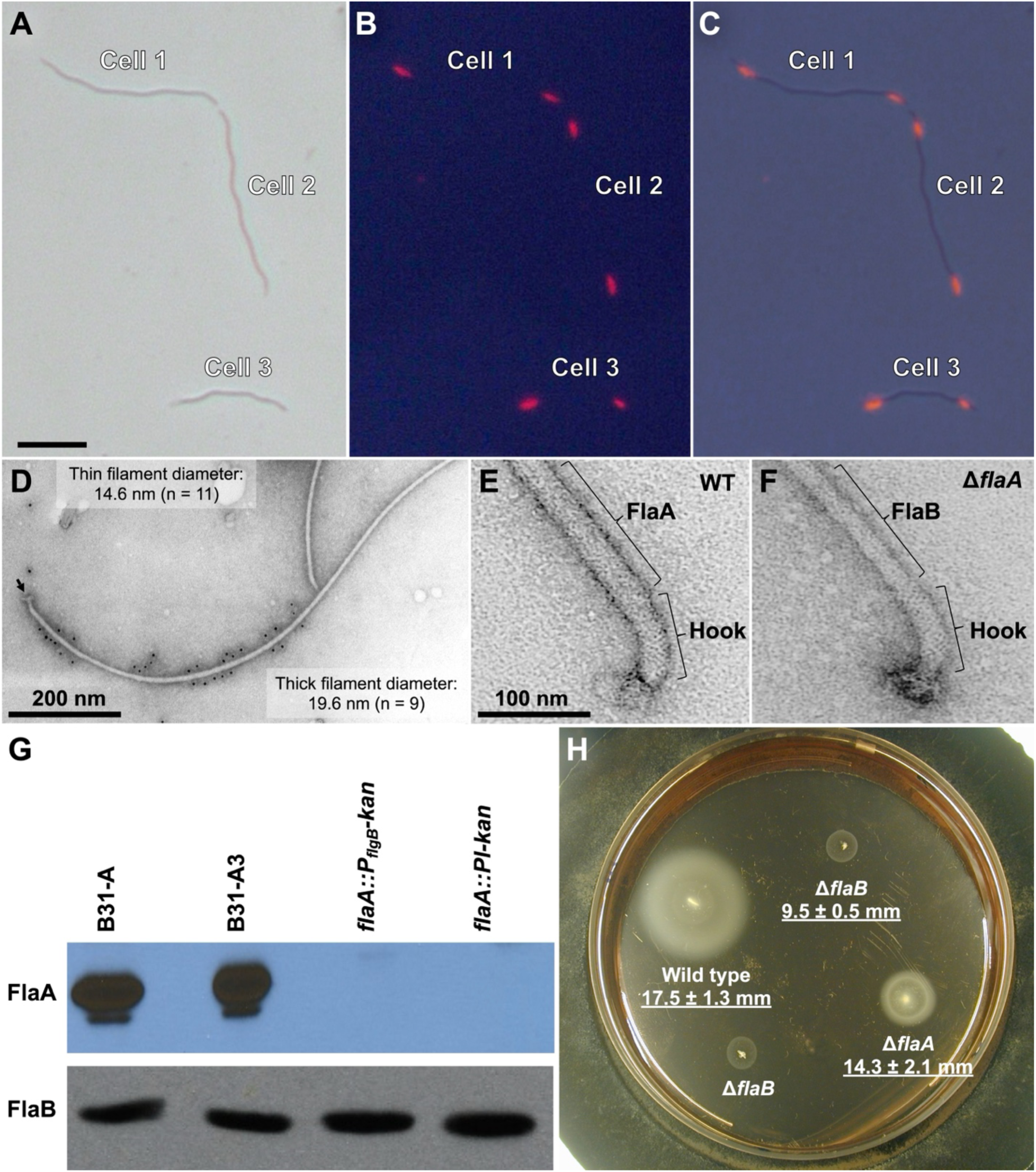
Characterization of flagellar proteins in *B. burgdorferi* and their roles in flagellar assembly and spirochetal motility. (**A–C**) Fluorescence microscopy of *B. burgdorferi* cells showing a phase-contrast image (A), immunofluorescence labeling with an anti-FlaA antibody (B), and the merged image (C). FlaA fluorescence is localized to the cell poles, corresponding to the proximal regions of the periplasmic flagella. (Scale bar, 5 µm.) (**D**) Negative-stain electron micrograph of purified periplasmic flagella showing two filament populations with distinct diameters. The mean diameters are 14.6 nm for thin filaments (n = 11) and 19.6 nm for thick filaments (n = 9). The arrow indicates the basal body. (Scale bar, 200 nm.) (**E** and **F**) Negative-stain electron micrographs of purified flagellar filaments from wild-type (E) and Δ*flaA* (F) cells. Thick filaments are absent in the Δ*flaA* mutant, indicating that FlaA is required for formation of the thick, sheathed filaments. (Scale bar, 100 nm, applies to both panels.) (**G**) Immunoblot analysis of whole-cell lysates from wild-type B31-A and B31-A3 strains, isogenic *flaA::PflgB-kan*, and *flaA::Pl-kan* strains using anti-FlaA (top) and anti-FlaB (bottom) antibodies. FlaA is undetectable in both *flaA* insertion mutants, whereas FlaB levels remain unchanged. (**H**) Swimming plate motility assay of wild-type, Δ*flaA*, and Δ*flaB* strains. Mean swimming diameters are 17.5 ± 1.3 mm for wild type, 14.3 ± 2.1 mm for Δ*flaA*, and 9.5 ± 0.5 mm for Δ*flaB*, indicating that loss of FlaA reduces motility, whereas loss of FlaB causes a more severe defect.

To determine whether this localized FlaA distribution corresponds to distinct filament architectures, we purified periplasmic flagella and examined them by immunogold electron microscopy using anti-FlaA antibodies. Filaments with two different diameters are readily visible (Fig. 1*D*). The thick filament diameter is similar to that observed previously (28). Immuno-gold particles specifically decorated the thick filaments but were absent from the thin filaments, showing that FlaA forms a sheath surrounding the FlaB filament core. To confirm this observation, we inactivated the *flaA* gene and verified the mutant by immunoblotting (Fig. 1*G*). Consistent with this observation, purified periplasmic flagella from the Δ*flaA* mutant contained only the thin-filament population, whereas the thick filaments observed in wild-type cells were completely absent (Fig. 1*E, F*). Immunoblotting with FlaB-specific antibodies further confirmed that FlaA absence did not affect synthesis of major filament protein FlaB (Fig. 1*G*). Thus, loss of the thick filament results directly from FlaA absence rather than from secondary effects on filament assembly. These results demonstrate that FlaA is required for assembly of the thick, sheathed filament near the flagellar motor.

We next examined the functional consequences of FlaA loss in motility by swimming plate assays. Compared with wild-type cells, the Δ*flaA* mutant exhibited markedly reduced swimming motility, and disruption of *flaB* completely abolished motility (Fig. 1*H*) (6, 9). These findings demonstrate that FlaA is dispensable for assembly of the FlaB filament core but is essential for sheath formation and contributes substantially to efficient spirochetal motility. By contrast, FlaB forms the filament core and is indispensable for flagellar function (6, 9).

Together, these biochemical, immunological, and functional analyses demonstrate that the *B. burgdorferi* periplasmic flagellum consists of a conserved FlaB core surrounded by a spatially restricted FlaA sheath assembled only over the proximal region adjacent to the flagellar motor (Fig. 2*A*).

**Fig. 2.**
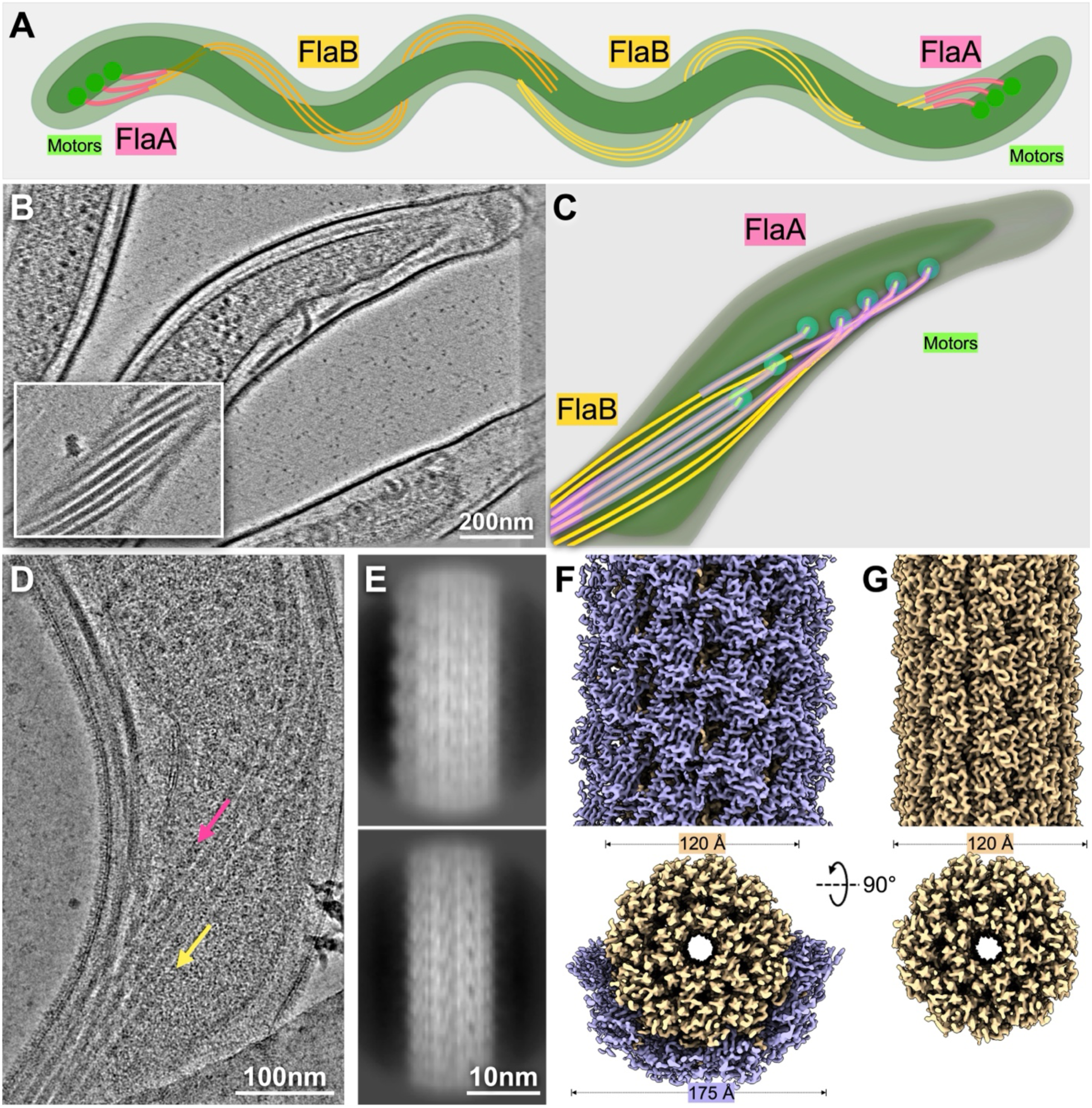
*In situ* structures of the *B. burgdorferi* periplasmic flagellar filament. (**A**) Schematic model of a *B. burgdorferi* cell showing bipolar periplasmic flagella anchored by flagellar motors at each cell pole. The flagella extend toward the cell center and wrap around the cell cylinder within the periplasmic space. The filament core consists of FlaB, whereas the FlaA sheath is predominantly localized to the pole-proximal regions. (**B**) Cryo-ET slice through a *B. burgdorferi* cell pole showing a bundle of periplasmic flagella between the inner and outer membranes. The boxed region highlights the filament bundle in a different tomographic slice. (**C**) Three-dimensional segmentation of the cell pole showing the cell cylinder (dark green), FlaB filament cores (yellow), FlaA sheath (Purple), and flagellar motors (light green). (**D**) Cryo-EM micrograph of a cell tip showing individual periplasmic flagella within the bundle. Arrows indicate representative thin (yellow) and thick (purple) filaments. (**E**) Representative 2D class averages of filament segments extracted from cryo-EM micrographs, showing sheathed (top) and unsheathed (bottom) filament classes. (**F** and **G**) Near-atomic structures of half-sheathed (F) and unsheathed (G) filaments, shown as surface representations viewed along the filament axis (top) and corresponding cross-sectional views (bottom). The FlaB core is colored yellow and the FlaA sheath purple. The FlaB core has a diameter of approximately 120 Å in both structures, whereas the FlaA sheath increases the overall filament diameter to approximately 175 Å where present. Cross-sectional views are oriented from the distal end, and all structures are shown at the same scale.

### *In situ* cryo-EM reveals an asymmetric *B. burgdorferi* periplasmic flagellar filament

To better understand the native architecture of the *B. burgdorferi* periplasmic flagellar filament, we combined cryo-electron tomography (cryo-ET) with *in situ* single-particle cryo-EM to visualize periplasmic flagella *in situ*. Cryo-ET analyses of wild-type cell poles showed that the FlaA sheath is predominantly associated with the pole-proximal portion of the filament, whereas the distal regions are composed primarily of the FlaB core (Fig. 2*B, C*; Movie S1). To define the structural basis of this spatial heterogeneity, we performed *in situ* single-particle cryo-EM analysis of the filaments in intact cells and resolved two major filament architectures: half-sheathed and non-sheathed filaments (Figs. 2*D–G*; *SI Appendix*, Fig. S2). The half-sheathed filament comprises of a FlaB core decorated by FlaA sheath, while the unsheathed filament consists entirely of FlaB (Fig. 2*F, G*). A less abundant class exhibited minor-sheathed FlaA coverage, likely representing an intermediate stage of sheath assembly (*SI Appendix*, Fig. S3). The FlaB core maintained a diameter of approximately 120 Å, whereas the FlaA assembly increased the overall filament diameter to approximately 175 Å (Fig. 2*F, G*). Importantly, FlaA assembles asymmetrically along one side of the FlaB core rather than forming an extensive sheath covering the FlaB core. This distinctive sheath architecture suggests that FlaA preferentially associates with the convex surface of the FlaB core, potentially contributing to filament curvature and supercoiling.

Cross-sectional views further revealed that the FlaB core comprises 11 protofilaments surrounding a central channel approximately 30 Å in diameter. In the half-sheathed reconstruction, six FlaA subunits were detected per helical repeat, although only the four central subunits were sufficiently resolved for atomic model building (*SI Appendix*, Fig. S5). The two peripheral subunits exhibited weaker, more diffuse densities, suggesting increased conformational flexibility or partial occupancy (Fig. 2*F*). Consistent with this observation, local resolution was highest within the FlaB core and decreased toward the outer surface of the FlaA sheath, indicating greater structural heterogeneity in the sheath (*SI Appendix*, Fig. S4). Nevertheless, well-resolved side-chain densities in the ordered regions of both FlaA and FlaB enabled unambiguous model building (Fig. 2*F*; *SI Appendix*, Fig. S5; *SI Appendix*, Table S1; Movie S2). In the minor-sheathed reconstruction, only three FlaA protofilaments exhibited discernible density, further highlighting the variable coverage and structural heterogeneity of the FlaA sheath (*SI Appendix*, Fig. S3).

These structural findings are consistent with our immunofluorescence and immunogold labeling analyses, which indicate that FlaA is predominantly localized to the pole-proximal regions of the periplasmic flagella (Fig. 1*A–D*). Therefore, these observations support a model in which FlaA sheath coverage progressively decreases along the filament, transitioning from a half-sheathed proximal region to an unsheathed distal region composed solely of FlaB.

### The FlaB filament core adopts a conserved spirochetal flagellin architecture

The near-atomic structure of the *B. burgdorferi* periplasmic flagellar filament enabled de novo atomic model building of the FlaB core (Fig. 3*A*). Like flagellins from other spirochetes (19, 29, 30), FlaB adopts a truncated flagellin fold comprising only the conserved D0 and D1 domains, lacking the surface-exposed D2 and D3 domains characteristic of enteric flagellins (16). The N-terminal (residues 1–32) and C-terminal (residues 300–336) segments form two long α-helices that associate to form the characteristic coiled-coil D0 domain, whereas the intervening sequence folds into a mixed α/β D1 domain (Fig. 3*B*). Within the filament, 11 FlaB protofilaments assemble into the characteristic five-start helical architecture (Fig. 3*A*). The D0 helices form the inner wall surrounding the central channel, and the D1 domains constitute the outer surface of the filament. Along each protofilament, adjacent FlaB subunits form an extensive longitudinal interface that is organized into two structural layers (Fig. 3*C, D*). Within the filament lumen, the N- and C-terminal helices of successive protomers interdigitate to form a continuous coiled-coil bundle extending along the filament axis. On the filament exterior, adjacent D1 domains pack head-to-tail, forming a continuous ridge along each protofilament.

**Fig. 3.**
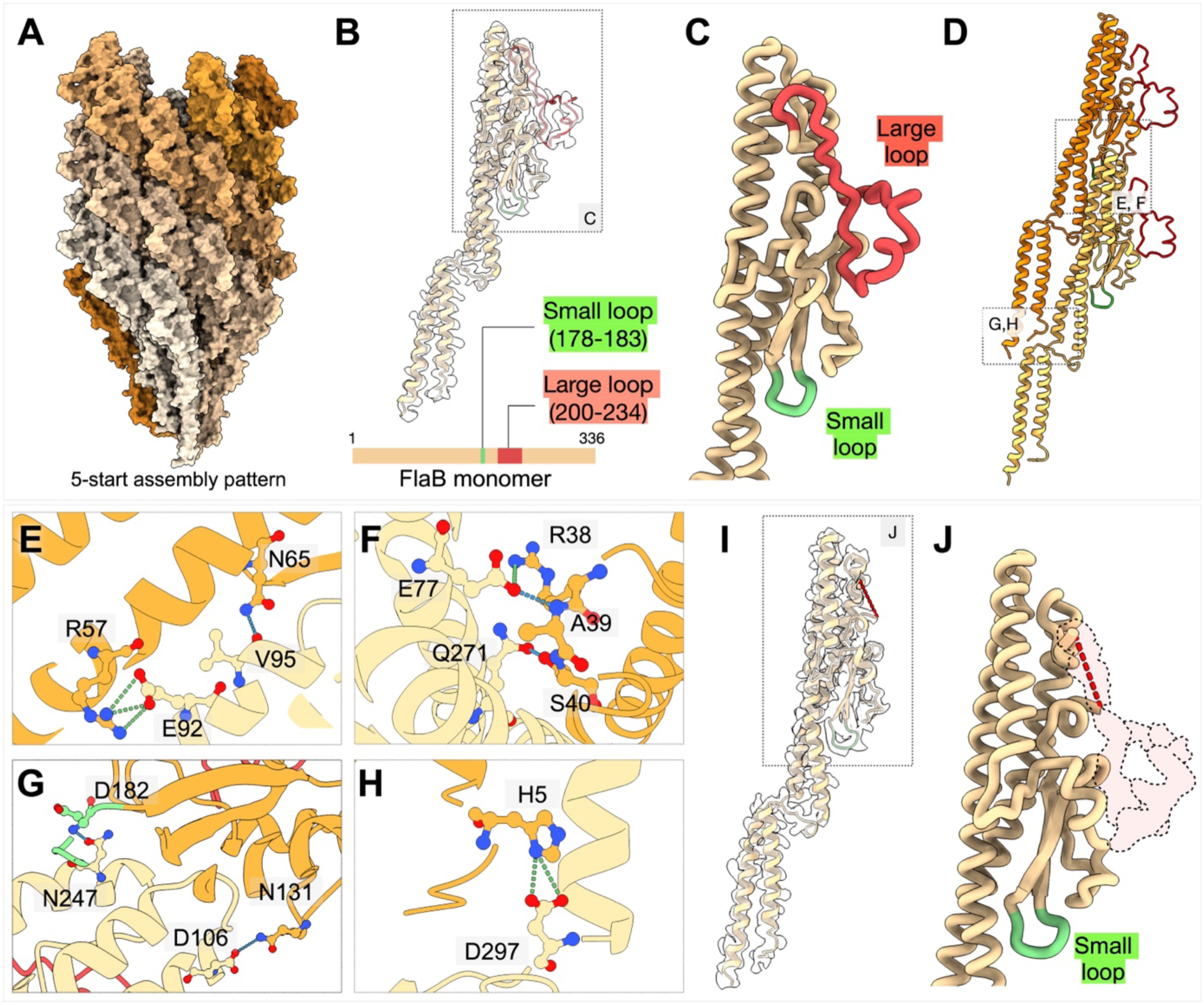
Atomic structure of the FlaB filament core. (**A**) Surface representation of the FlaB filament core viewed along the filament axis, illustrating its five-start helical architecture. Individual protofilaments are shown in alternating shades to distinguish neighboring helical strands. (**B**) Ribbon representation of a FlaB protomer associated with FlaA (not shown). Two surface-exposed loops are highlighted in red (large loop) and green (small loop). Their positions are indicated in the schematic of the 336-residue FlaB polypeptide below. The large loop comprises residues 200–234, whereas the small loop comprises residues 178–183. (**C**) Enlarged view of the outer surface of the FlaB protomer highlighting the large and small loops. (**D**) Ribbon representation of two adjacent FlaB subunits within a single protofilament. (**E–H**) Close-up views of longitudinal interactions between neighboring FlaB subunits. The upper and lower subunits are colored orange and pale yellow, respectively. Hydrogen bonds and salt bridges are indicated by blue and green dashed lines, respectively. The interaction networks involve R57-E92 and N65-V95 (E), R38-E77, A39-E77, and S40-Q271 (F), D182-N247 and N131-D106 (G), and H5-D297 (H). (**I**) Ribbon representation of a FlaB protomer lacking FlaA association. The large loop is unresolved in the absence of FlaA. (**J**) Enlarged view of the boxed region in I, highlighting the unresolved large loop.

The longitudinal interface is further stabilized by an extensive network of intermolecular interactions (Fig. 3*E–H*). Three salt bridges (R57–E92, R38–E77, and H5–D297), together with multiple hydrogen bonds involving N65–V95, A39–E77, S40–Q271, D182–N247, and N131– D106, reinforce the interactions between successive FlaB protomers, with residues from the upper subunit listed first in each pair. These interactions establish a continuous axial scaffold that maintains the structural integrity and mechanical stability of the filament core.

Notably, two prominent surface loops, designated the large loop (residues 200–234) and small loop (residues 178–183), extend from the D1 domain toward the filament exterior (Fig. 3*B, C*). The large loop of FlaB was well resolved in protomers associated with FlaA but became largely unresolved in those lacking bound FlaA (Fig. 3*I, J*). By contrast, the small loop remained well resolved regardless of FlaA association. These observations suggest that FlaA binding selectively stabilizes the large loop of FlaB, promoting its local ordering and potentially strengthening the interaction between the FlaB core and FlaA sheath.

### FlaA is anchored to FlaB through the large loop

To elucidate the molecular basis of sheath assembly, we built atomic models of FlaA and its interface with the FlaB filament core through the half-sheathed reconstruction (Fig. 4*A–C*). FlaA adopts a compact globular fold composed primarily of β-sheets flanked by several α-helices (Fig. 4*D*). Within the sheath, adjacent FlaA protomers form longitudinal arrays that follow the underlying FlaB protofilaments (Fig. 4*E*). Neighboring FlaA subunits interact through a relatively small interface stabilized by hydrogen bonds involving K109, Y174, D278, K204, and N231, together with hydrophobic interactions involving W206 and V229 (Fig. 4*F*). These interactions likely maintain sheath integrity while providing sufficient flexibility to accommodate the helical geometry and curvature of the filament.

**Fig. 4.**
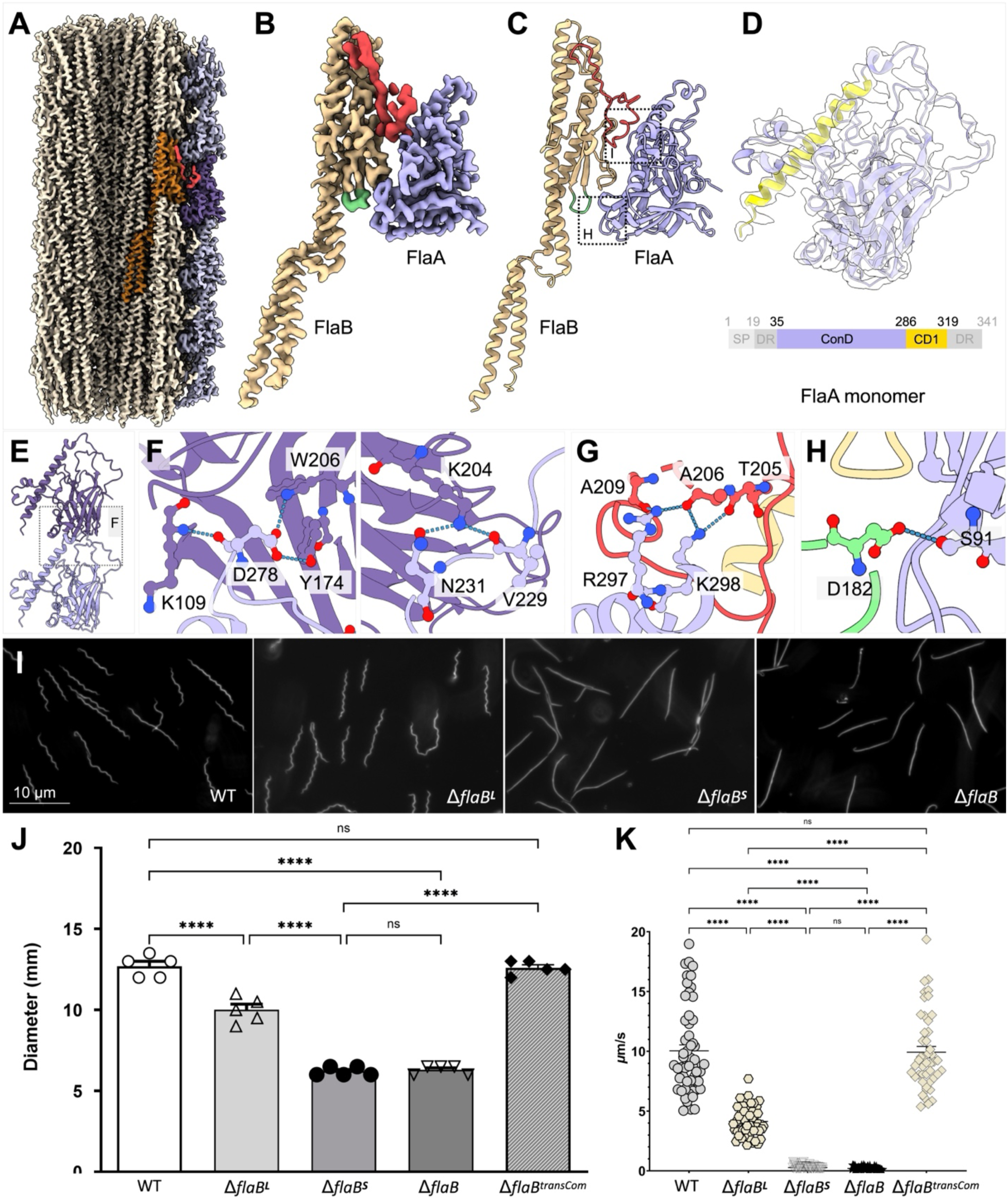
The FlaA sheath is anchored to the FlaB filament core. (**A**) Cutaway view of the segmented cryo-EM map of a half-sheathed filament showing the FlaB core (pale yellow) and FlaA sheath (purple). One FlaB protomer is highlighted in orange, with its large loop colored red, to illustrate its interaction with a neighboring FlaA protomer (purple). (**B**) Cryo-EM density map of a representative FlaB–FlaA complex. FlaB is colored yellow, with the large and small loops highlighted in red and green, respectively, and FlaA in purple. (**C**) Atomic model of the FlaB–FlaA complex, colored as in B. (**D**) Ribbon representation of a FlaA protomer. (**E**) Ribbon representation of two adjacent FlaA subunits within a single protofilament. (**F**) Close-up views of longitudinal interactions between neighboring FlaA subunits. Hydrogen bonds are indicated by blue dashed lines. The interaction networks involve K10-D278, Y174-D278, and W206 (top), and K204-N231 and V229 (bottom). (**G**) Close-up view of the FlaB–FlaA interface mediated by the distal portion of the FlaB large loop. FlaB residues T205, A206, and A209 interact with FlaA residues R297 and K298. Hydrogen bonds are indicated by blue dashed lines. (**H**) Close-up view of the FlaB–FlaA interface mediated by the FlaB small loop, highlighting the interaction between FlaB residue D182 and FlaA residue S91. (**I**) Representative dark-field or phase-contrast micrographs of wild-type (WT), Δ*flaB^L^*, Δ*flaB^S^*, and Δ*flaB* cells grown to mid-exponential phase in BSK-II medium at 34 °C. WT and Δ*flaB^L^* cells retain the characteristic flat-wave spirochetal morphology, whereas Δ*flaB^S^* and Δ*flaB* cells exhibit shorter, irregularly bent morphologies with substantial loss of the regular planar waveform. (**J**) Swimming plate motility assay of the indicated strains, including the trans-complemented Δ*flaB^transCom^* strain. Cells were inoculated into semisolid agarose plates containing BSK-II medium and incubated at 34 °C under 3.4% CO_2_ for 4-5 days before swimming diameters were measured. The Δ*flaB^L^* mutant exhibits a modest but significant reduction in swimming diameter relative to WT, whereas Δ*flaB^S^* and Δ*flaB* mutants show more severe, statistically indistinguishable motility defects, indicating that the FlaB small loop is particularly important for efficient motility. Motility is restored to WT levels in the Δ*flaB^transCom^*strain. Each symbol represents an independent biological replicate (n = 5 per strain). Bars indicate means ± SEM. Statistical significance was determined by one-way ANOVA followed by Tukey’s multiple-comparisons test (****P < 0.0001; ns, not significant). (**K**) Tracking analysis of the indicated strains.

Each FlaA protomer binds a single FlaB subunit primarily through the large loop and small loop of the D1 domain (Fig. 4*C, G, H*). These loops engage the inner surface of FlaA, forming the principal contact interface between the two proteins (Fig. 4*G, H*). The interface is stabilized by hydrogen bonds involving FlaA residues R297 and K298 and FlaB residues T205, A206, and A209 within the large loop, together with an additional interaction between S91of FlaA and D182 of FlaB within the small loop (Fig. 4*G, H*). Thus, FlaA attachment is mediated primarily by surface-exposed loops of FlaB, with the large loop providing the principal anchoring interface.

Consistent with this structural arrangement, the large loop adopts a well-defined conformation in FlaB protomers bound by FlaA (Fig. 3*C, D*) but becomes largely disordered in the absence of the FlaA sheath (Fig. 3*I, J*). This observation suggests that FlaA binding stabilizes an intrinsically flexible region of the FlaB core surface. To further characterize the molecular interface, we analyzed its electrostatic and hydrophobic properties. Electrostatic potential mapping revealed that the outer surface of the assembled filament is predominantly electronegative, whereas the FlaB surface exposed upon sheath removal is comparatively neutral, with localized patches of positive potential (*SI Appendix*, Fig. S6*A*). Hydrophobicity mapping showed that the FlaA surface contacting FlaB is more hydrophobic than its solvent-exposed outer surface, whereas the corresponding FlaB interface exhibits hydrophobicity comparable to the surrounding core surface (*SI Appendix*, Fig. S6*B*). Each FlaA protomer contacts the underlying FlaB core through a spatially contiguous interface involving 25 FlaB residues and 26 FlaA residues (*SI Appendix*, Fig. S7*A, B*). Interface analysis revealed buried solvent-accessible surface areas of 811.4 Å^2^ for FlaB and 763.3 Å^2^ for FlaA, representing less than 5% of the total solvent-accessible surface area of each protomer (*SI Appendix*, Fig. S7*C*). Collectively, these structural features suggest that FlaA is anchored to the FlaB core through surface loops.

To assess the functional significance of these interactions, we independently replaced the large and small loops with flexible glycine–serine linkers that preserve polypeptide continuity while removing their native interaction surfaces. Residues 200–234 of the large loop were replaced with (GGGGS)_2_, and residues 178–183 of the small loop were replaced with GGGGS. The resulting Δ*flaB^L^* and Δ*flaB^S^*strains were characterized alongside a Δ*flaB* mutant (*SI Appendix*, Fig. S8). Immunoblotting confirmed the expected protein expression profiles of the mutant strains (*SI Appendix*, Fig. S8*A*). Dark-field microscopy of mid-exponential-phase cultures revealed distinct effects of the two loop replacements on cell morphology (Fig. 4*I*). Δ*flaB^L^* cells retained the characteristic flat-wave form. By contrast, Δ*flaB^S^* cells exhibited rod-shaped morphologies. These morphological abnormalities closely resembled those of Δ*flaB* cells (6), suggesting that the small loop contributes more substantially than the large loop to maintaining normal cell morphology.

The morphological defects were accompanied by corresponding reductions in translational motility. In swimming plate assays, wild-type cells produced swimming rings measuring 15.25 ± 0.48 mm, whereas Δ*flaB^L^* cells exhibited a modest but significant reduction to 9.64 ± 0.24 mm (P < 0.0001; Fig. 4*J*). By contrast, Δ*flaB^S^* and Δ*flaB* cells displayed substantially smaller rings of 6.63 ± 0.38 mm. Consistent with these results, single-cell tracking analysis showed that Δ*flaB^S^* and Δ*flaB* cells were nonmotile (Movie S3, S4), whereas Δ*flaB^L^* cells retained motility (4.13 ± 0.17 μm/s; Movie S5), albeit less efficiently than wild-type cells (10.04 ± 0.50 μm/s) (Fig. 4*J*; Movie S6). Trans-complementation restored wild-type swimming behavior, confirming that the observed defects resulted from the engineered alterations at the *flaB* locus (Fig. 4*J*; Movie S7).

The more severe phenotype of Δ*flaB^S^* is consistent with the structural location of the small loop. Unlike the large loop, the small loop lies at the longitudinal interface between adjacent FlaB subunits within a protofilament (Fig. 3*D*). In particular, D182 contributes to the interactions between successive FlaB protomers (Fig. 3*G*). Replacing the small loop may therefore disrupt longitudinal subunit packing of FlaB as well as weaken FlaA binding, explaining the pronounced effects of the small loop on cell morphology and motility.

### Asymmetric FlaA sheath amplifies supercoiling of the *B. burgdorferi* periplasmic flagellum

To determine how asymmetric sheath assembly influences filament geometry, we compared the structures of unsheathed and half-sheathed regions of the native supercoiled periplasmic flagellum (Fig. 5). Both regions retained the conserved 11-protofilament FlaB core, whereas the half-sheathed region was decorated by an asymmetric FlaA sheath restricted to the convex face of the supercoil (Fig. 5*A*). This organization suggests that differences in sheath coverage arise primarily from variations in FlaA assembly.

**Fig. 5.**
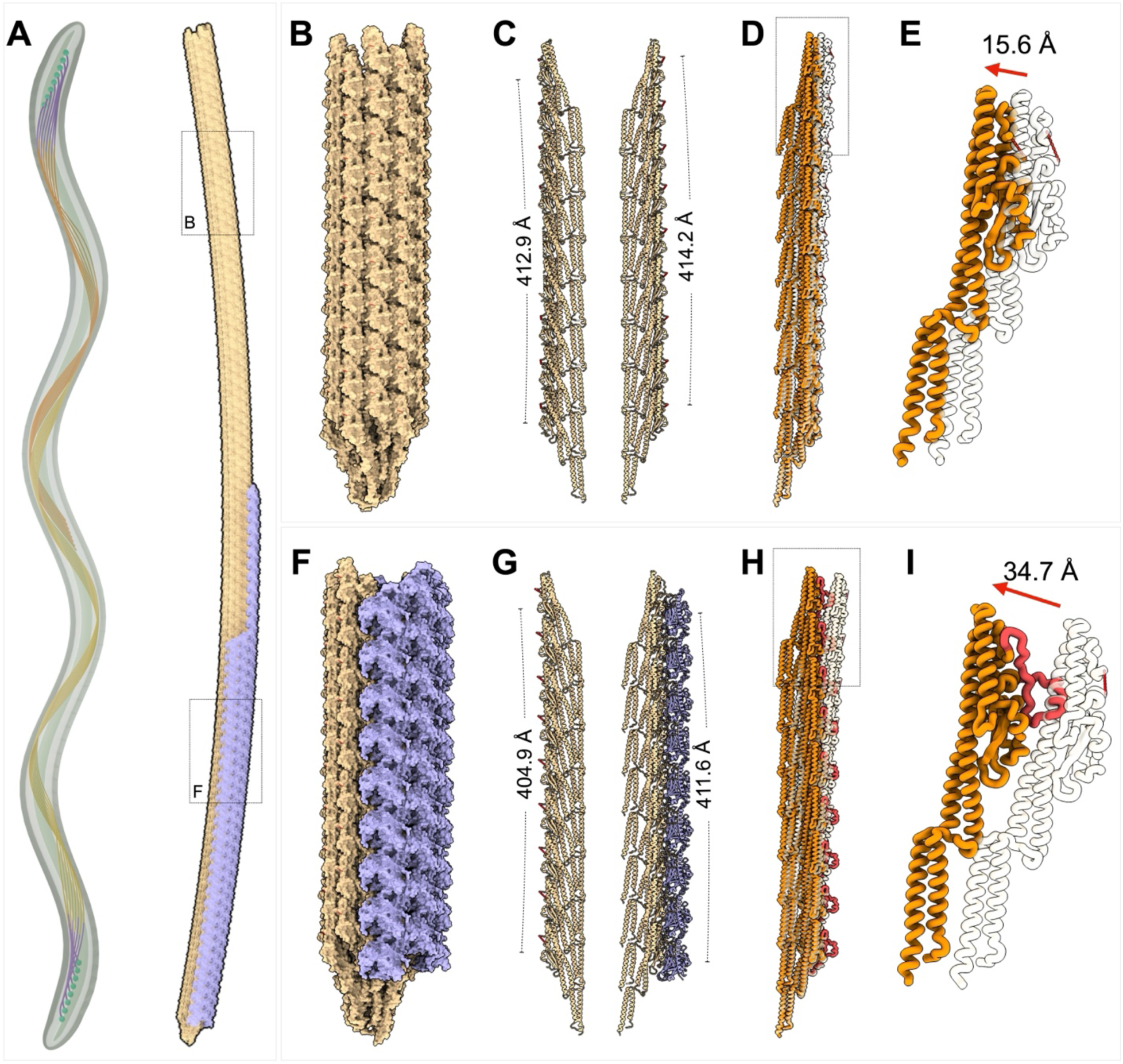
Asymmetric architecture of the supercoiled *B. burgdorferi* periplasmic flagellar filament. (**A**) Overview of a *B. burgdorferi* cell and its periplasmic flagellar bundle. A representative supercoiled filament is shown on the right, with the FlaB core colored yellow and the FlaA sheath purple. The filament contains a continuous transition between unsheathed and half-sheathed segments. The boxed regions are enlarged in the upper and lower rows. (**B–E**) Structural features of the unsheathed filament segment. (**B**) Surface representation of the FlaB filament core lacking the FlaA sheath. (**C**) Comparison of protofilaments on the concave (left) and convex (right) sides of the supercoil. The axial distances spanning nine helical repeats are 412.9 Å and 414.2 Å on the concave and convex sides, respectively, corresponding to a difference of 1.3 Å. (**D**) Lateral displacement across the filament segment, with the red arrow indicating an offset of 8.4 Å. (**E**) Ribbon representation of the same region, with surface-exposed loops omitted for clarity. The lateral offset becomes apparent beginning at the fourth helical repeat. (**F–I**) Structural features of the half-sheathed filament segment, displayed as in B–E. (**F**) Surface representation of the cryo-EM reconstruction showing the half-sheathed filament. (**G**) Comparison of protofilaments on the concave (left) and convex (right) sides. The axial distances spanning nine helical repeats are 404.9 Å and 411.6 Å, respectively, corresponding to a difference of 6.7 Å. (**H**) Lateral displacement across the half-sheathed filament segment, with the red arrow indicating an offset of 34.7 Å. (**I**) Ribbon representation of a FlaB protomer from the half-sheathed filament, with the large loop highlighted in red. Axial distances in C and G were measured between equivalent atoms in subunits n and n + 9 along the same protofilament. The concave and convex sides are defined relative to the supercoil axis shown in A.

The unsheathed filament exhibited relatively modest geometric asymmetry between the concave and convex sides of the supercoil (Fig. 5*B–E*). Over nine successive FlaB subunits, the axial distance measured 412.9 Å on the concave side and 414.2 Å on the convex side, corresponding to a longitudinal register difference of only 1.3 Å (Fig. 5*C*). This small axial asymmetry was accompanied by a lateral displacement of 15.6 Å (Fig. 5*D, E*), indicating that the FlaB core alone adopts a slightly curved supercoiled conformation.

By contrast, the half-sheathed filament exhibited substantially greater geometric asymmetry (Fig. 5*F–I*). The corresponding axial distances measured 404.9 Å on the concave side and 411.6 Å on the convex side, yielding a longitudinal register difference of 6.7 Å (Fig. 5*G*). The lateral displacement also increased markedly, from 15.6 Å in the unsheathed filament to 34.7 Å in the half-sheathed filament (Fig. 5*H, I*). In both structures, these differences accumulated progressively along the filament, becoming more pronounced after approximately the fourth helical repeat.

Despite these pronounced differences in filament geometry, individual FlaB protomers retained essentially the same conformation, indicating that supercoiling is associated primarily with subtle changes in intersubunit packing rather than substantial conformational rearrangements within individual protomers. Importantly, the unsheathed filament retained measurable longitudinal asymmetry and a supercoiled conformation, demonstrating that curvature is an intrinsic property of the FlaB core. The asymmetric FlaA sheath therefore does not generate supercoiling but instead amplifies the intrinsic curvature of the filament by enhancing the geometric asymmetry between its concave and convex sides.

Collectively, these findings reveal that the *B. burgdorferi* periplasmic flagellar filament exhibits pronounced structural asymmetry that is further enhanced by selective FlaA assembly on the convex face of the supercoil near the cell poles (Movie S2). By increasing longitudinal asymmetry without substantially altering the conserved FlaB core architecture, the FlaA sheath may modulate filament curvature and mechanical properties, providing a structural basis for understanding how sheath assembly contributes to the distinctive morphology and motility of spirochetes.

## Discussion

Periplasmic flagella are defining features of spirochetes and are essential for their distinctive morphology, motility, and ability to navigate complex environments (5, 18, 31). Recent structural studies of *Leptospira* and *Treponema* have revealed that their periplasmic filaments consist of a conserved FlaB core surrounded by an asymmetric, multiprotein sheath (19, 29, 30), establishing an architecture fundamentally distinct from that of the external flagellar filaments of *Salmonella* (16) and *Vibrio* (32). Despite these advances, the molecular organization of the periplasmic flagellar filament in the Lyme disease spirochete *B. burgdorferi*, a major model for investigating spirochetal motility and pathogenesis, has remained poorly understood (6, 9, 21). Here, using *in situ* single-particle cryo-EM, we determined near-atomic structures of native *B. burgdorferi* periplasmic flagellar filament, revealing striking asymmetry in both sheath architecture and its spatial distribution along the filament. Although the FlaB core retains the conserved 11-protofilament organization, a single sheath protein, FlaA, assembles asymmetrically along its convex surface, predominantly within the proximal filament region near the cell poles. This spatially restricted sheath organization distinguishes *B. burgdorferi* from other characterized spirochetes and reveals a distinct mechanism for locally modulating filament architecture and potentially its mechanical properties.

Our structures reveal how FlaA modulates filament geometry without substantially altering the conserved FlaB core. FlaA binds primarily to a large surface-exposed loop unique to *B. burgdorferi* FlaB, stabilizing the otherwise flexible region while preserving the underlying filament architecture. Comparison of half-sheathed and unsheathed filaments demonstrates that FlaB possesses intrinsic curvature and supercoiling, which are further amplified by asymmetric FlaA assembly along the convex surface. Similar observations in *Leptospira* and *Treponema* suggest that intrinsic curvature is a conserved feature of spirochetal flagella, whereas species-specific sheath proteins provide distinct mechanisms for modulating asymmetric filament architecture and curvature (19, 29, 30).

A striking feature of the *B. burgdorferi* filament is the preferential localization of FlaA to the pole-proximal region, where motor-generated torque is transmitted to the filament through the hook. This spatial organization suggests that FlaA provide localized mechanical reinforcement in a region subjected to substantial torsional and bending loads during flagellar rotation. By increasing filament diameter and introducing additional intermolecular interactions, the FlaA sheath enhance mechanical stability while preserving flexibility along the distal filament. Consistent with its functional importance, deletion of *flaA* or replacement of the large surface-exposed loop of FlaB impairs motility (Fig. 1*H*, 4*J*, *K*). Together, these observations suggest that spatially restricted FlaA assembly provide a mechanism for balancing proximal reinforcement with distal flexibility, thereby optimizing filament mechanics for efficient motility (Movie S8).

In summary, our study reveals a previously unrecognized mechanism linking asymmetric protein assembly to filament curvature, the characteristic flat-wave morphology, and motility of the Lyme disease spirochete. More broadly, our findings uncover a distinct architectural strategy for modulating flagellar geometry and mechanical properties, providing new insights into the molecular basis of spirochetal motility and pathogenesis. Future studies will be needed to elucidate how sheath assembly is spatially regulated and how asymmetric sheath formation influences filament mechanics and torque transmission.

## Materials and Methods

### Bacterial strains, culture conditions, and oligonucleotide primers

High-passage *Borreliella burgdorferi* sensu stricto strain B31A (wild-type) and its derivative mutants were cultured in Barbour-Stoenner-Kelly II (BSK-II) liquid medium or on semi-solid agar plates at 34°C with 3.4% CO_2_ as previously described (4, 13). Antibiotics selection was applied when necessary: kanamycin (300 μg/ml), gentamicin (40 μg/ml). *Escherichia coli* NEB 5-alpha strain (New England Biolabs, Carlsbad, CA, USA) was used for plasmid cloning and amplification. The antibiotic concentrations used for E. coli selection were as follows: kanamycin 50 μg/ml, gentamicin 5 μg/ml, and ampicillin 100 μg/ml.

### Construction of *flaB* (bb0147) in-frame deletion mutant and trans-complementation

To generate an in-frame deletion of *flaB* (gene locus, *BB0147*), targeted gene disruption constructs were generated using a PCR-based overlap extension method as described previously (13). Briefly, primers P_3_/P_4_ and P_5_/P_6_-containing sequence overlaps complementary to the kanamycin resistance cassette (*aph1*) were designed to amplify approximate 1-kb regions flanking the *flaB* coding sequence. Following initial amplification of the individual 5’- and 3’-flanking and *aph1* fragments, a fusion PCR was performed using primers P_3_/P_6_ to join all three fragments, generating the *flaBBb::aph1* disruption construct. Mutant clones were selected on semi-solid BSK-II agar plates containing kanamycin (300 μg/ml).

To construct a trans-complementation vector, the *flaB* native promoter and open reading frame (ORF) were PCR-amplified using primer pair P_7_/P_8_. The resulting PflaB-flaB fragment was cloned into the pJET1.2/blunt cloning vector (Thermo Fisher Scientific, Waltham, MA, USA), excised using BamHI and PstI and subcloned into corresponding sites of the *B. burgdorferi* shuttle vector pBSV2G to generate the complementation plasmid PflaB-flaB/pBSV2G. The resulting contract was transformed into *flaB* mutant cells, and transformants were selected on semi-solid BSK-II agar plates containing kanamycin (300 μg/ml) and gentamicin (40 μg/ml). All oligonucleotide primers utilized in this study are listed in *SI Appendix*, Table S2.

### Construction of *flaB* truncation mutants

Two *flaB* genes containing engineered loop replacements were synthesized by GenScript (Piscataway, NJ, USA). The large loop (^200^GEGAQTAQAAPVQEGVQQEGAQQPAPATAPS QGGV^234^) was replaced with a flexible (GGGGS)₂ linker, whereas the small loop(^178^GANQDE^183^) was replaced with a (GGGGS) linker. Both constructs were driven by the native *flaB* promoter and flanked by engineered 5′-BamHI and 3′-PstI restriction sites. The synthesized fragments were digested with BamHI and PstI and ligated into the corresponding sites of the *B. burgdorferi* shuttle vector pBSV2G, generating the trans-complementation plasmids pBSV2G-L2 (large-loop replacement) and pBSV2G-S1 (small-loop replacement). The resulting constructs were transformed into the *flaB* deletion mutant using the same protocol as described above.

### Bacterial motion tracking analysis, swim plate assays

Swimming velocity of *B. burgdorferi* cells was evaluated using a computer-based motion tracking system as previously described (33). Briefly, active motile, logarithmic-phase cultures were diluted 1:1 in BSKII medium and then 20 μl of diluted cultures were mixed with an equal volume of 2% methylcellulose ((viscosity 4,000 cP; MC4000). Cell movement was recorded using iMovie software on an Apple Mac computer. Videos are exported as QuickTime files and imported into OpenLab (Improvision Inc., Coventry, UK) for frame cropping, calibration, and conversion to LIFF format. Individual trajectories and velocities were quantified using Volocity software (Improvision Inc., Coventry, UK). For each strain, at least 20 individual cells were tracked for up to 30 sec.

Swimming plate assays was performed using 0.35% agarose plates prepared with BSK-II medium diluted 1:10 in Dulbecco’s phosphate-buffered saline (dPBS, pH 7.5) as previously described (4, 6, 13). In brief, *B. burgdorferi* wild type, the *flaB* deletion mutant, and three trans-complemented strains with the full-length *flaB* or two truncated versions were inoculated in soft agarose plates and incubated for 4–5 days at 34°C under 3.4% CO_2_. Swimming ring diameters were documented in millimeters (mm), and average values were determined from four independent plates.

### Gel electrophoresis and immunoblot analysis

Sodium dodecyl sulfate-polyacrylamide gel electrophoresis (SDS-PAGE) and enhanced chemiluminescence (ECL) immunoblotting were performed as described previously (13). Briefly, whole-cell lysates (10-20 μg protein per lane) were separated on 12%SDS-PAGE gels and transferred onto polyvinylidene difluoride (PVDF) membrane (Bio-Rad Laboratories, Hercules, CA, USA). Membranes were blocked and probed with specific primary antibodies. Monoclonal and polyclonal antibodies were generously provided by the following investigators: monoclonal anti-FlaB (H9724) by A. G. Barbour (University of California, Irvine, CA), monoclonal anti-DnaK by J. Benach (SUNY at Stony Brook, NY). Polyclonal antibodies against FlaA were prepared as previously reported (34). Blots were incubated with horseradish peroxidase (HRP)-conjugated secondary antibodies and developed using an ECL luminol substrate. Chemiluminescent signals were captured using a ChemiDoc MP imaging system and quantified using Image Lab software (Bio-Rad Laboratories). All immunoblot analyses were performed using three independent biological replicates.

### Construction of a *B. burgdorferi flaA* mutant

The *flaA* gene (*bb0668*) was inactivated using a promoterless Pl-Kan cassette, as described (6). The Pl-Kan cassette contains the ribosomal binding site and transcription start site of the *B. burgdorferi flgB* promoter (PflgB), followed by the kanamycin resistance gene (35). PCR amplification of *flaA*, construction of the inactivation plasmids, and electroporation of linear *flaA*-Pl-Kan DNA into competent B31A cells were performed as described previously (6, 36). Briefly, *flaA* DNA was PCR-amplified using primers P9 and P10 and then ligated into the pGEM-T Easy vector (Promega Inc.), yielding pflaA-Easy. Pl-Kan was similarly PCR-amplified from PflgB-Kan with primers P11 and P12, which contain HindIII restriction sites. The HindIII-restricted Pl-Kan DNA was then inserted into the HindIII sites within *flaA*, yielding pflaA-Pl-Kan. This deletion– insertion was in-frame. Restriction mapping indicated that the direction of transcription of Pl-Kan was the same as that of *flaA*. The pflaA-Pl-Kan plasmid was linearized using NotI, and the purified DNA was electroporated into competent *B. burgdorferi* cells. Mutants were selected on solid growth medium containing 200 μg/ml kanamycin. The kanamycin-resistant clones were verified for *flaA* inactivation by immunoblotting using *B. burgdorferi*-specific anti-FlaA polyclonal antisera. All oligonucleotide primers used in this study are listed in *SI Appendix*, Table S2.

### Immunofluorescence microscopy and immunoelectron microscopy

Immunofluorescence microscopy was conducted to localize FlaA as previously described (37). Briefly, 1.5 ml log-phase *B. burgdorferi* cultures were harvested, washed twice with PBS buffer (pH 7.5), and then treated with methanol at −20°C for 1 h. The collected cells were then treated with lysozyme (1 mg ml-1) in GTE buffer (50 mM glucose, 25 mM Tris, and 1 mM EDTA, pH 7.5) for 1 h at room temperature. Cells were then placed on poly-L-lysine-coated coverslips and allowed to fully air dry. The obtained coverslips were first incubated in a blocking solution (2% BSA in PBS, pH 7.5) for 1 h, followed by incubation in the blocking solution containing1:500 diluted FlaA antibody for 1 h at room temperature. Finally, the coverslips were washed five times with PBS, incubated with the secondary goat anti-mouse Texas red antibody (Invitrogen) for 1 h at room temperature, washed with PBS, and mounted in 40% glycerol for image processing as described previously (37).

Immunogold labeling of purified periplasmic flagella was performed using an anti-FlaA antibody. The flagella were purified using Triton X-100, as described previously (14, 20, 21, 38). About 5 μl of the purified flagellar suspension was combined with 5 μl of the FlaA antibody solution, diluted in 10 mM Tris, 5 mM EDTA, 150 mM NaCl (pH 8.0). This mixture was incubated on ice for 20 min. Next, add 2 μl of 5-nm colloidal gold-conjugated anti-rabbit antibody, diluted in 10 mM Tris, 5 mM EDTA, 300 mM NaCl (pH 8.0), and incubate at room temperature for 1 hour. The mixture was placed on a hydrophilic-treated carbon-coated copper grid, washed three times with 10 mM Tris-HCl, 5 mM EDTA (pH 8.0), negatively stained with 2% phosphotungstic acid (pH 7.0), and observed with a JEOL 1020 microscope at 80 kV.

The purified periplasmic flagella were negatively stained with 2% phosphotungstic acid (pH 7.0) on carbon-collodion-coated copper grids and observed with a JEOL 1020 microscope, as described above.

### Cryo-EM sample preparation

For *in situ* cryo-EM, *B. burgdorferi* cells were collected by centrifugation at 2,000 ×g for approximately 5 min in 1.5-mL microcentrifuge tubes. The cell pellet was gently washed once with phosphate-buffered saline (PBS) and resuspended in PBS to a final optical density of OD_600_ = 1.0 before plunge freezing. 5 μL of bacterial culture was applied to freshly glow-discharged Quantifoil R2/1 200-mesh copper grids (Quantifoil Micro Tools GmbH). Grids were blotted with Whatman filter paper for approximately 6 s and rapidly plunge-frozen into a liquid ethane–propane mixture using a GP2 plunger (Leica Microsystems). During vitrification, the environmental chamber of the GP2 was maintained at 25°C and 90% relative humidity.

### *In situ* cryo-EM data collection and processing

Cryo-EM data were collected on a Titan Krios transmission electron microscope (Thermo Fisher Scientific) operated at 300 kV and equipped with a field-emission gun, a BioQuantum energy filter (20-eV slit width), and a K3 Summit direct electron detector (Gatan). Images were acquired in nanoprobe mode at a nominal magnification of 81,000× using SerialEM (39) with a customized multishot acquisition script that specifically targeted cell tips on cryo-EM grids. Imaging conditions included a spot size of 5, a C2 aperture of 50 or 70 μm, an objective aperture of 100 μm, and defocus values ranging from −1.6 to −2.2 μm. Movies were recorded in super-resolution mode with a total accumulated dose of approximately 70 e⁻/Å^2^. During Patch Motion Correction in cryoSPARC, an F-crop factor of 2 was applied, resulting in a calibrated pixel size of 1.068 Å. In total, 39,784 micrographs were collected from wild-type *B. burgdorferi B31* cells.

Movie frames were motion-corrected using Patch Motion Correction (40), and contrast transfer function (CTF) parameters were estimated with Patch CTF Estimation (41) in cryoSPARC (42). Filament segments were automatically picked using the Filament Tracer module (43) with a filament diameter of 200 Å, a minimum inter-particle spacing of 100 Å, and a minimum filament length of 400 Å. Particles were extracted with a box size of 360 pixels. After reference-free two-dimensional classification to remove contaminants and poorly aligned particles, 608,922 filament segments were retained for subsequent three-dimensional reconstruction.

An initial model was generated by *ab initio* reconstruction and further refined by heterogeneous refinement followed by local refinement. Helical symmetry parameters were determined using the Symmetry Search Utility (44) in cryoSPARC, yielding an optimal solution with a helical rise of 4.70 Å and a twist of 65.40°. Application of these parameters produced a helical reconstruction of the sheathed flagellar filament at an overall resolution of 3.60 Å.

To resolve structural heterogeneity within the filament sheath, 3D classification was performed using a sheath-focused mask. This analysis separated the dataset into three structural classes corresponding to half-sheathed, minor-sheathed, and non-sheathed filaments. Particles from each class were subjected to iterative local refinement, during which the rotational search range was gradually reduced from 20° to 2°, the translational search range from 10 Å to 2 Å, and the initial low-pass filter from 12 Å to 3 Å. The final reconstructions of sheathed and non-sheathed filament contained 198,388 particles and 330,787 particles (after particle expansion), reaching Fourier shell correlation (FSC, 0.143 criterion) resolutions of 3.41 Å and 3.20 Å, respectively. The additional class of minor-sheathed filament contained 30,823 particles, reaching Fourier shell correlation (FSC, 0.143 criterion) resolutions of 4.07 Å.

To determine the spatial distribution of the FlaA sheath along native periplasmic flagella, particles were re-extracted using a larger box size of 1,280 pixels and Fourier-binned to 448 pixels to preserve the local curvature of the filament and its surrounding cellular context. Following two-dimensional classification, classes exhibiting both pronounced filament curvature and adjacent inner membrane density were selected for homogeneous reconstruction. The resulting low-resolution 3D reconstruction visualized by UCSF ChimeraX (45) revealed that the FlaA sheath is consistently positioned on the convex side of the supercoiled filament, opposite the cell cylinder. Finally, refined particles were mapped back onto the original micrographs to verify their locations within intact cells and confirm the native spatial distribution of the sheath along individual periplasmic flagella.

### Model building and structural analysis

The amino acid sequences of FlaB and FlaA were obtained from UniProt (FlaB, accession no. P11089; FlaA, accession no. P70856) and used to generate initial structural models with AlphaFold3 (46). The predicted models were rigid-body fitted into the cryo-EM density maps using UCSF ChimeraX (45), followed by iterative manual rebuilding in Coot (47) and real-space refinement in Phenix (48). The final atomic models showed excellent stereochemical quality based on the validation statistics reported by Phenix (48). Model-to-map correlation coefficients were 0.78 for the half-sheathed filament, and 0.79 for the non-sheathed filament, indicating good agreement between the atomic models and the corresponding cryo-EM reconstructions. Several surface-exposed loop regions, particularly BLoop3 in the absence of the FlaA sheath, exhibited weak or discontinuous density and were either modeled with reduced confidence or omitted, consistent with their intrinsic conformational flexibility.

Cryo-EM map visualization, segmentation, molecular surface rendering, and figure preparation were performed using UCSF ChimeraX (45). Protein–protein interfaces and intermolecular interactions were analyzed using PDBe PISA (49).

### Cryo-ET data collection and analysis

Frozen-hydrated *B. burgdorferi* cells were imaged on a 300-kV Titan Krios transmission electron microscope (Thermo Fisher Scientific). Tilt series were acquired using SerialEM (39) with a bidirectional tilt scheme spanning −48° to +48° in 3° increments, yielding 33 projection images per tilt series. Data were collected at a nominal defocus of −1 μm with a cumulative electron dose of approximately 70 e⁻/Å^2^ distributed evenly across all tilts.

Movie frames were corrected for beam-induced motion using MotionCor2(40), followed by tilt-series alignment and stack generation in IMOD (50). Tomograms were reconstructed from 4× Fourier-binned tilt series using Tomo3D (51). The 3D organization of the cell envelope and periplasmic flagella was visualized by manual segmentation using Dragonfly (version 2022.2; Comet Technologies Inc., Quebec, Canada). The outer membrane, inner membrane, flagellar motors, and periplasmic flagellar filaments were manually segmented based on tomographic densities. Regions corresponding to the FlaA sheath were additionally segmented to visualize their spatial distribution relative to the filament core, flagellar motors, and cell envelope.

## Acknowledgments

The project was supported by grants R01AI087946, R01AI132818, and R01AI078958 from the National Institute of Allergy and Infectious Diseases (NIAID), R01DE023080 and R01DE034063 from the National Institute of Dental, Craniofacial Research (NIDCR). Cryo-ET data were collected at Yale CryoEM resource, which is funded in part by the NIH grant 1S10OD023603-01A1. We thank the Yale Center for Research Computing facility (YCRC) for providing computing support and advice and Jennifer Aronson for critical reading and editing of the manuscript.

## Reference

1. N. W. Charon, S. F. Goldstein, Genetics of motility and chemotaxis of a fascinating group of bacteria: the spirochetes. Annu Rev Genet 36, 47–73 (2002).

2. M. Strnad et al., Nanomechanical mechanisms of Lyme disease spirochete motility enhancement in extracellular matrix. Commun Biol 4, 268 (2021).

3. J. D. Radolf, M. J. Caimano, B. Stevenson, L. T. Hu, Of ticks, mice and men: understanding the dual-host lifestyle of Lyme disease spirochaetes. Nat Rev Microbiol 10, 87–99 (2012).

4. C. Li et al., Asymmetrical flagellar rotation in *Borrelia burgdorferi* nonchemotactic mutants. Proc Natl Acad Sci U S A 99, 6169–6174 (2002).

5. S. Nakamura, Spirochete Flagella and Motility. Biomolecules 10 (2020).

6. M. A. Motaleb et al., *Borrelia burgdorferi* periplasmic flagella have both skeletal and motility functions. Proc Natl Acad Sci U S A 97, 10899–10904 (2000).

7. M. Harman, D. K. Vig, J. D. Radolf, C. W. Wolgemuth, Viscous dynamics of Lyme disease and syphilis spirochetes reveal flagellar torque and drag. Biophys J 105, 2273–2280 (2013).

8. K. Abe, T. Kuribayashi, K. Takabe, S. Nakamura, Implications of back-and-forth motion and powerful propulsion for spirochetal invasion. Sci Rep 10, 13937 (2020).

9. S. Z. Sultan et al., Motility is crucial for the infectious life cycle of *Borrelia burgdorferi*. Infect Immun 81, 2012–2021 (2013).

10. M. Zamba-Campero et al., Broadly conserved FlgV controls flagellar assembly and *Borrelia burgdorferi* dissemination in mice. Nat Commun 15, 10417 (2024).

11. S. Z. Sultan et al., Motor rotation is essential for the formation of the periplasmic flagellar ribbon, cellular morphology, and *Borrelia burgdorferi* persistence within Ixodes scapularis tick and murine hosts. Infect Immun 83, 1765–1777 (2015).

12. K. H. Moon et al., Spirochetes flagellar collar protein FlbB has astounding effects in orientation of periplasmic flagella, bacterial shape, motility, and assembly of motors in *Borrelia burgdorferi*. Mol Microbiol 102, 336–348 (2016).

13. K. Zhang et al., FlhF regulates the number and configuration of periplasmic flagella in *Borrelia burgdorferi*. Mol Microbiol 113, 1122–1139 (2020).

14. Y. Ge, C. Li, L. Corum, C. A. Slaughter, N. W. Charon, Structure and expression of the FlaA periplasmic flagellar protein of *Borrelia burgdorferi*. J Bacteriol 180, 2418–2425 (1998).

15. Y. Ge, N. W. Charon, An unexpected flaA homolog is present and expressed in *Borrelia burgdorferi*. J Bacteriol 179, 552–556 (1997).

16. K. Yonekura, S. Maki-Yonekura, K. Namba, Complete atomic model of the bacterial flagellar filament by electron cryomicroscopy. Nature 424, 643–650 (2003).

17. J. L. Fields et al., Structural diversity and clustering of bacterial flagellar outer domains. Nat Commun 15, 9500 (2024).

18. F. San Martin, L. Fule, G. Iraola, A. Buschiazzo, M. Picardeau, Diving into the complexity of the spirochetal endoflagellum. Trends Microbiol 31, 294–307 (2023).

19. J. Wang et al., Asymmetric architecture and adaptation of *Treponema* flagella. Nat Commun 10.1038/s41467-026-74267-7 (2026).

20. M. S. Sal et al., Borrelia burgdorferi uniquely regulates its motility genes and has an intricate flagellar hook-basal body structure. J Bacteriol 190, 1912–1921 (2008).

21. M. A. Motaleb, M. S. Sal, N. W. Charon, The decrease in FlaA observed in a flaB mutant of *Borrelia burgdorferi* occurs posttranscriptionally. J Bacteriol 186, 3703–3711 (2004).

22. E. A. Wunder et al., FcpB Is a Surface Filament Protein of the Endoflagellum Required for the Motility of the Spirochete. Front Cell Infect Microbiol 8, 130 (2018).

23. E. A. Wunder et al., A novel flagellar sheath protein, FcpA, determines filament coiling, translational motility and virulence for the Leptospira spirochete. Mol Microbiol 101, 457–470 (2016).

24. K. Kurniyati, Y. Chang, J. Liu, C. Li, Transcriptional and functional characterizations of multiple flagellin genes in spirochetes. Mol Microbiol 118, 175–190 (2022).

25. C. Li et al., The spirochete FlaA periplasmic flagellar sheath protein impacts flagellar helicity. J Bacteriol 182, 6698–6706 (2000).

26. C. Li, C. W. Wolgemuth, M. Marko, D. G. Morgan, N. W. Charon, Genetic analysis of spirochete flagellin proteins and their involvement in motility, filament assembly, and flagellar morphology. J Bacteriol 190, 5607–5615 (2008).

27. J. Liu et al., Intact flagellar motor of *Borrelia burgdorferi* revealed by cryo-electron tomography: evidence for stator ring curvature and rotor/C-ring assembly flexion. J Bacteriol 191, 5026–5036 (2009).

28. C. Dombrowski et al., The elastic basis for the shape of *Borrelia burgdorferi*. Biophys J 96, 4409–4417 (2009).

29. A. Kawamoto, T. Kuribayashi, M. Morita, S. Nakamura, N. Koizumi, Asymmetric sheath coordination controls flagellar architecture and function in *Leptospira* spirochete. EMBO J 45, 2882–2904 (2026).

30. F. San Martin et al., Core-sheath coupling controls flagellar curvature and motility in *Leptospira*. Nat Commun 17 (2026).

31. N. W. Charon et al., The unique paradigm of spirochete motility and chemotaxis. Annu Rev Microbiol 66, 349–370 (2012).

32. W. Guo et al., Structures of the sheathed flagellum reveal mechanisms of assembly and rotation in *Vibrio cholerae*. Nat Microbiol 10, 3305–3314 (2025).

33. R. G. Bakker, C. Li, M. R. Miller, C. Cunningham, N. W. Charon, Identification of specific chemoattractants and genetic complementation of a *Borrelia burgdorferi* chemotaxis mutant: flow cytometry-based capillary tube chemotaxis assay. Appl Environ Microbiol 73, 1180–1188 (2007).

34. K. Zhang, C. W. Sze, H. Zhao, J. Liu, C. Li, *Borrelia burgdorferi* serine protease HtrA is a pleiotropic regulator of stress response, motility, flagellar hemostasis, and infectivity. Commun Biol 8, 341 (2025).

35. J. L. Bono et al., Efficient targeted mutagenesis in *Borrelia burgdorferi*. J Bacteriol 182, 2445–2452 (2000).

36. M. A. Motaleb, M. R. Miller, R. G. Bakker, C. Li, N. W. Charon, Isolation and characterization of chemotaxis mutants of the Lyme disease Spirochete *Borrelia burgdorferi* using allelic exchange mutagenesis, flow cytometry, and cell tracking. Methods Enzymol 422, 421–437 (2007).

37. C. Li, H. Xu, K. Zhang, F. T. Liang, Inactivation of a putative flagellar motor switch protein FliG1 prevents *Borrelia burgdorferi* from swimming in highly viscous media and blocks its infectivity. Mol Microbiol 75, 1563–1576 (2010).

38. K. Zhang et al., Two CheW coupling proteins are essential in a chemosensory pathway of *Borrelia burgdorferi*. Mol Microbiol 85, 782–794 (2012).

39. D. N. Mastronarde, Automated electron microscope tomography using robust prediction of specimen movements. J Struct Biol 152, 36–51 (2005).

40. S. Q. Zheng et al., MotionCor2: anisotropic correction of beam-induced motion for improved cryo-electron microscopy. Nat Methods 14, 331–332 (2017).

41. A. J. Noble et al., Routine single particle CryoEM sample and grid characterization by tomography. Elife 7 (2018).

42. A. Punjani, J. L. Rubinstein, D. J. Fleet, M. A. Brubaker, cryoSPARC: algorithms for rapid unsupervised cryo-EM structure determination. Nat Methods 14, 290–296 (2017).

43. S. T. Huber, T. Kuhm, C. Sachse, Automated tracing of helical assemblies from electron cryo-micrographs. J Struct Biol 202, 1–12 (2018).

44. E. H. Egelman, A robust algorithm for the reconstruction of helical filaments using single-particle methods. Ultramicroscopy 85, 225–234 (2000).

45. T. D. Goddard et al., UCSF ChimeraX: Meeting modern challenges in visualization and analysis. Protein Sci 27, 14–25 (2018).

46. J. Abramson et al., Accurate structure prediction of biomolecular interactions with AlphaFold 3. Nature 10.1038/s41586-024-07487-w (2024).

47. P. Emsley, K. Cowtan, Coot: model-building tools for molecular graphics. Acta Crystallogr D Biol Crystallogr 60, 2126–2132 (2004).

48. D. Liebschner et al., Macromolecular structure determination using X-rays, neutrons and electrons: recent developments in Phenix. Acta Crystallogr D Struct Biol 75, 861–877 (2019).

49. E. Krissinel, K. Henrick, Inference of macromolecular assemblies from crystalline state. J Mol Biol 372, 774–797 (2007).

50. J. R. Kremer, D. N. Mastronarde, J. R. McIntosh, Computer visualization of three-dimensional image data using IMOD. J Struct Biol 116, 71–76 (1996).

51. J. I. Agulleiro, J. J. Fernandez, Tomo3D 2.0--exploitation of advanced vector extensions (AVX) for 3D reconstruction. J Struct Biol 189, 147–152 (2015).

