## Supplemental data for "Asymmetric flagellar filaments shape *Borreliella burgdorferi* morphology and motility"

**Supplemental Information**

A

| Name | Locus tag | Length | MW (kDa) | Isoelectronic point | Function | Signal peptides |
|---|---|---|---|---|---|---|
| FlaB | BB_0147 | 336 aa | 35.7 | 5.37 | Core | No |
| FlaA | BB_0668 | 341 aa | 38.4 | 7.69 | Sheath | Yes |

B

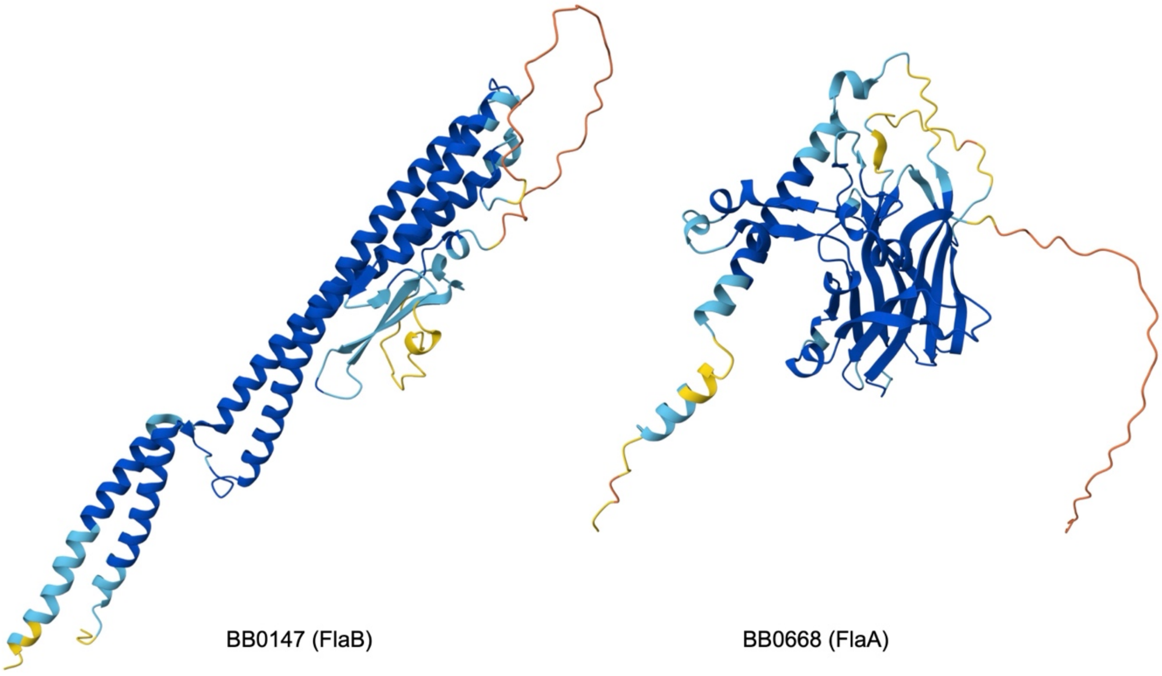


***SI Appendix*, Fig. S1. Sequence features and predicted structures of the two flagellar filament proteins in *B. burgdorferi*.** (**A**) Summary of the sequence features of FlaB (BB_0147) and FlaA (BB_0668), including locus tag, polypeptide length, molecular weight, theoretical isoelectric point (pI), localization within the filament, and presence of a predicted signal peptide. Molecular weights and theoretical pI values were calculated using ExPASy ProtParam. (**B**) Ribbon representations of AlphaFold2-predicted structures of FlaB (left) and FlaA (right), retrieved from the AlphaFold Protein Structure Database (accession numbers AF-P11089-F1 for FlaB and AF-P70856-F1 for FlaA). Structures are colored according to predicted local distance difference test (pLDDT) scores, ranging from dark blue (high confidence) to orange (low confidence).

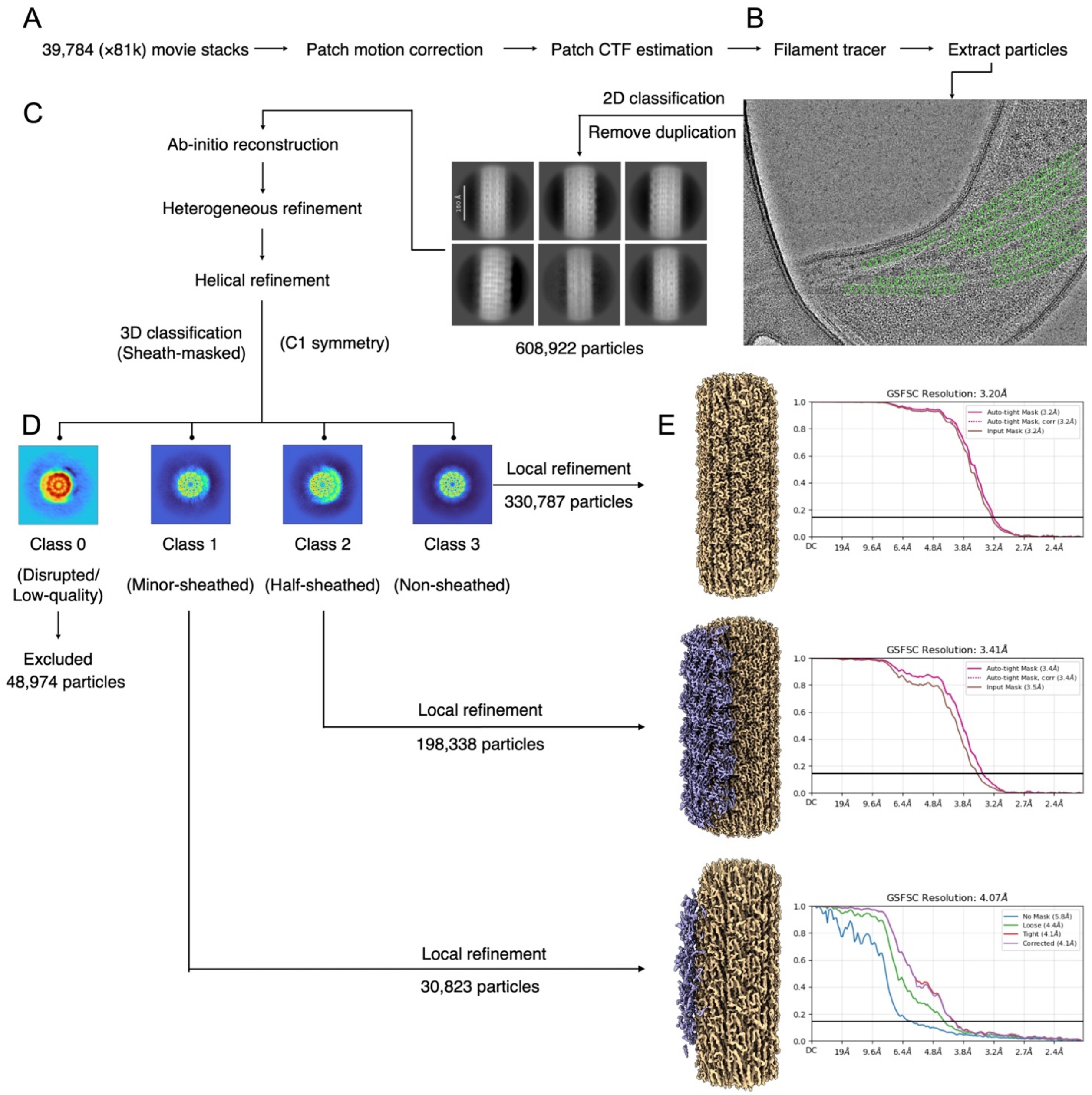


**SI Appendix, Fig. S2. Cryo-EM data processing workflow for the *B. burgdorferi* periplasmic flagellar filament.** (**A**) A total of 39,784 movie stacks collected at a nominal magnification of ×81,000 were processed using patch motion correction and patch CTF estimation in cryoSPARC v5.0.6. (**B**) Filaments were identified using the filament tracer, and particles were extracted with a box size of 360 pixels. A representative micrograph is shown with traced filaments highlighted in green. (**C**) Following 2D classification and duplicate removal, 608,922 particles were retained for *ab initio* reconstruction, heterogeneous refinement, and helical refinement with C1 symmetry. (Scale bar in 2D class averages, 100 Å.) (**D**) Sheath-masked 3D classification resolved four distinct classes. Class 0 (disrupted or low-quality filaments, 48,974 particles) was excluded from further analysis. Class 1 (minor-sheathed, 30,823 particles), Class 2 (half-sheathed, 198,338 particles), and Class 3 (unsheathed, 330,787 particles) were subjected to separate local refinements. (**E**) Final cryo-EM density maps and corresponding gold-standard Fourier shell correlation (FSC) curves for the three filament classes, with resolutions of 4.07 Å, 3.41 Å, and 3.20 Å, respectively, determined at the FSC = 0.143 criterion. The FlaB filament core is colored beige and the FlaA sheath purple.

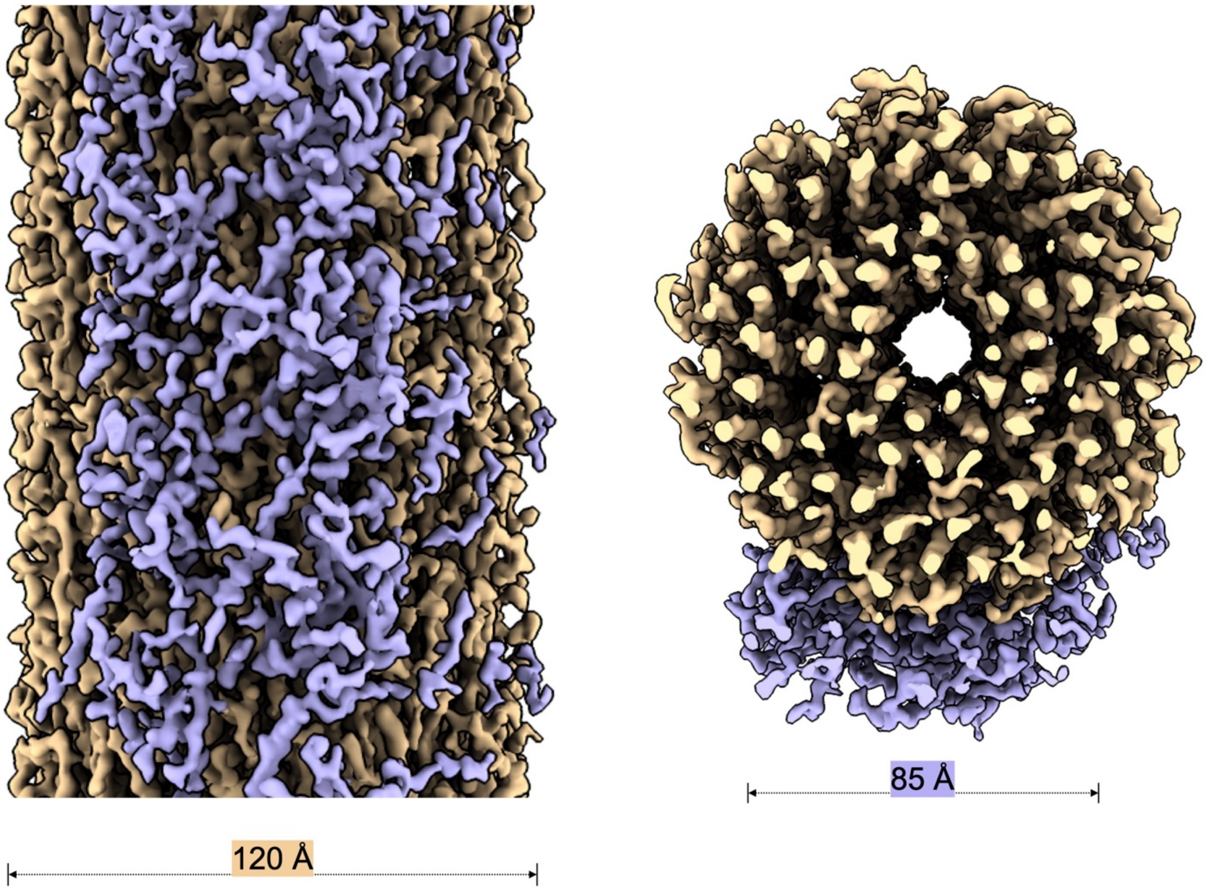


***SI Appendix*, Fig. S3. Cryo-EM reconstruction of the minor-sheathed periplasmic flagellar filament.** (Left) Side view of the cryo-EM density map showing the FlaB core (beige) and FlaA sheath (purple). The FlaA sheath forms a narrow longitudinal band along the filament rather than extending around its entire circumference. (Right) Cross-sectional view along the filament axis showing that the FlaA sheath occupies a limited arc along the outer surface of the FlaB core. The FlaB core has an outer diameter of approximately 120 Å, whereas the sheath-covered region spans approximately 85 Å in width.

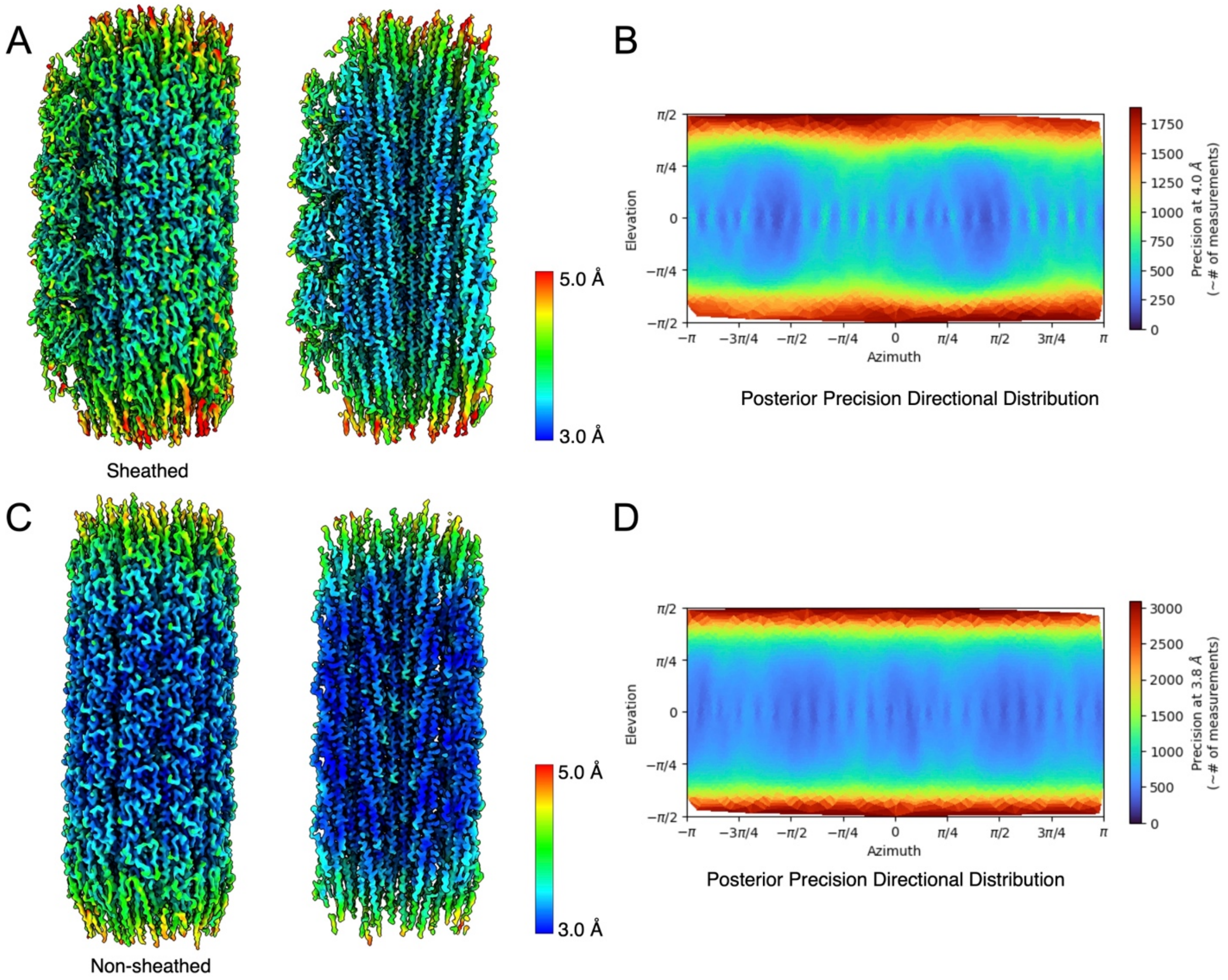


***SI Appendix*, Fig. S4. Local resolution and directional resolution estimates of the periplasmic flagellar filament reconstructions.** (**A** and **B**) Half-sheathed filament reconstruction. (**C** and **D**) Unsheathed filament reconstruction. (**A** and **C**) Cryo-EM density maps colored according to local resolution, as estimated in cryoSPARC (42). Each reconstruction is shown as an outer surface view (left) and a longitudinal cutaway view (right). (**B** and **D**) Posterior precision directional distributions showing the number of measurements contributing to each viewing direction at resolutions of 4.0 Å for the half-sheathed filament (B) and 3.8 Å for the unsheathed filament (D).

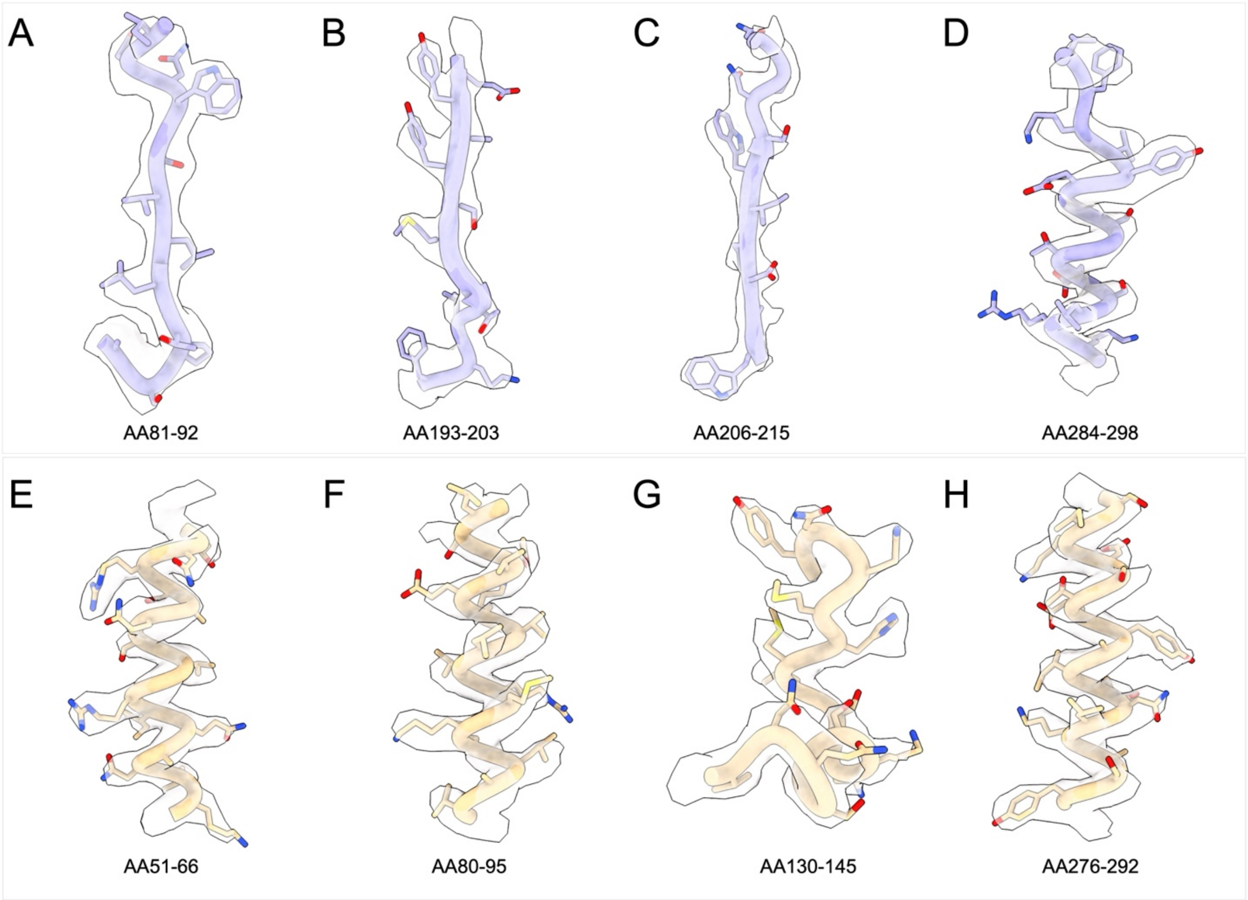


***SI Appendix*, Fig. S5. Representative cryo-EM densities of the half-sheathed filament with fitted atomic models.** (**A–D**) Representative regions of FlaA corresponding to residues 81–92 (A), 193–203 (B), 206–215 (C), and 284–298 (D). (**E–H**) Representative regions of FlaB corresponding to residues 51–66 (E), 80–95 (F), 130–145 (G), and 276–292 (H). Cryo-EM density maps are shown as surface representations, with fitted atomic models displayed as sticks and the polypeptide backbones traced as ribbons.

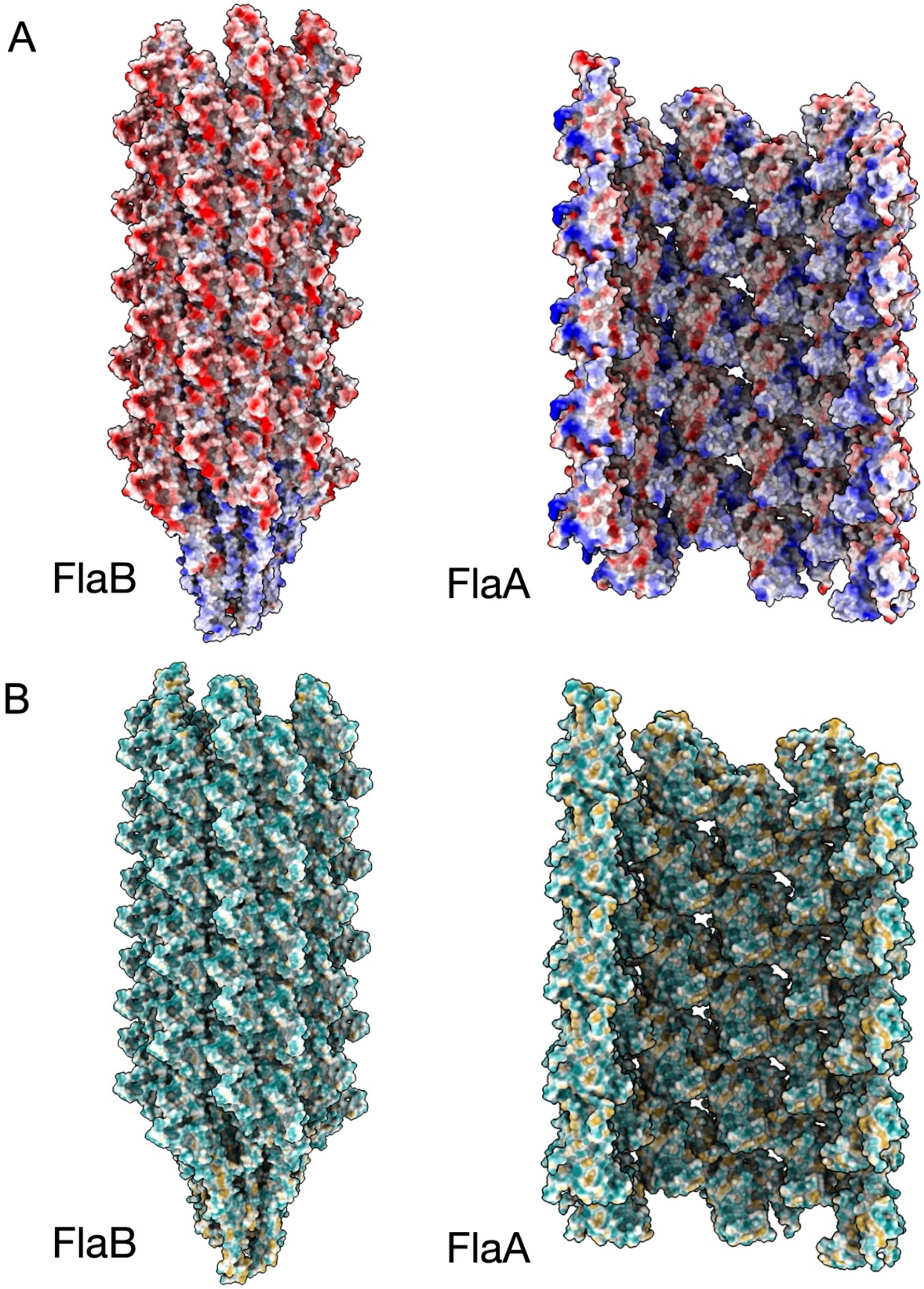


***SI Appendix*, Fig. S6. Surface properties of the *B. burgdorferi* periplasmic flagellar filament.** (**A**) Coulombic electrostatic potential mapped onto the molecular surfaces of the FlaB filament core (left) and FlaA sheath (right), viewed along the filament axis. Electrostatic potentials are colored from −10 to +10 kcal/(mol·e), with red indicating negative potential and blue indicating positive potential. (**B**) Molecular lipophilicity potential mapped onto the same molecular surfaces and shown in the same orientations as in A. Surface colors range from dark cyan (hydrophilic) to gold (hydrophobic).

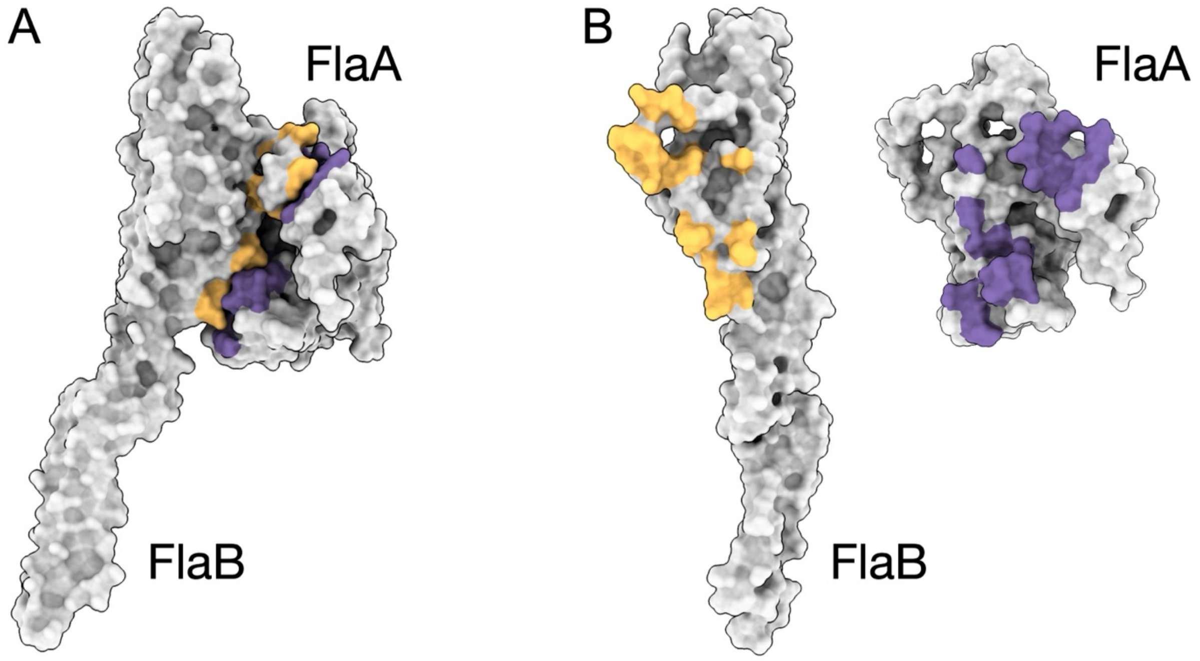


C

| Protein name | Interface residue number | Interface residue percentage | Interface area | Interface area percentage |
|---|---|---|---|---|
| FlaB | 25 | 7.4% | 811 Å$^2$ | 4.3% |
| FlaA | 26 | 9.0% | 763.3 Å$^2$ | 4.7% |

***SI Appendix*, Fig. S7. Molecular interactions between FlaA and FlaB in the periplasmic flagellar filament.** (**A**) Surface representations of a FlaB–FlaA protomer pair within the assembled filament, with interface residues highlighted in yellow on FlaB and purple on FlaA. (**B**) Open-book representation of the FlaB–FlaA interface, showing the two protomers separated and rotated to expose their interacting surfaces. (**C**) Analysis of the FlaB–FlaA interface using PDBePISA (49). The interface involves 25 residues of FlaB (7.4% of the polypeptide) and 26 residues of FlaA (9.0%), burying 811.4 Å$^2$ and 763.3 Å$^2$ of solvent-accessible surface area, respectively. These buried areas correspond to 4.3% and 4.7% of the total solvent-accessible surface areas of FlaB and FlaA, respectively.

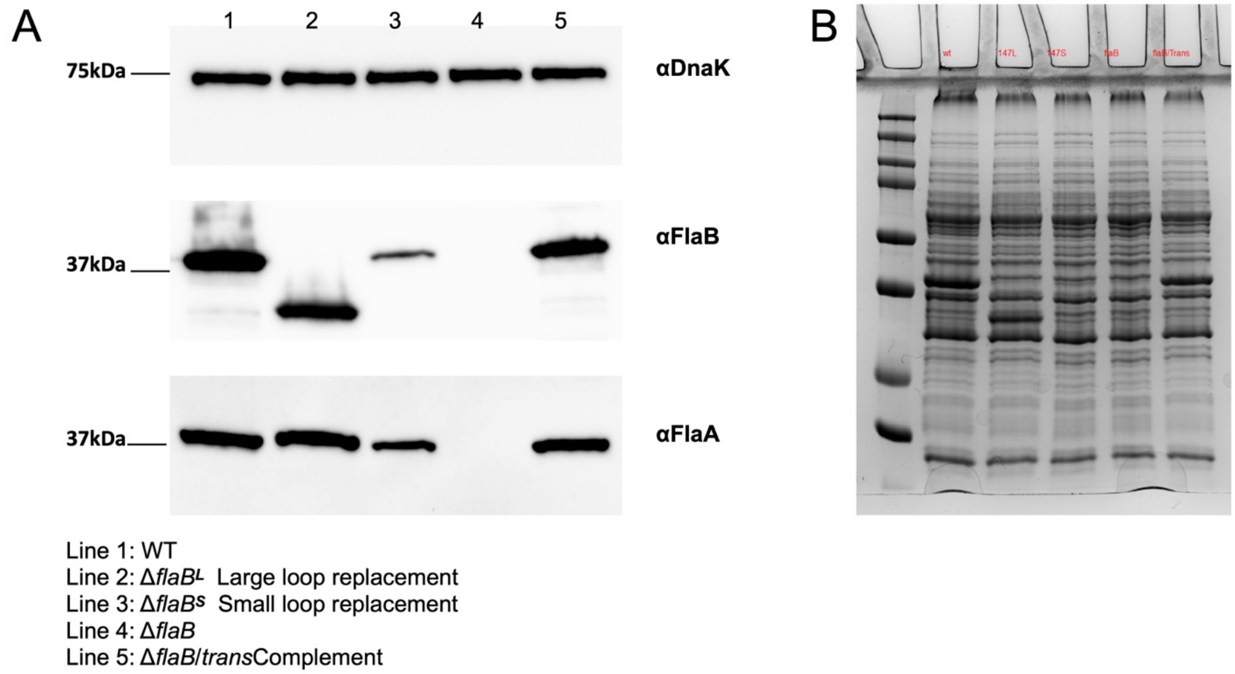


***SI Appendix*, Fig. S8. Immunoblotting analysis of *flaB* mutants.** Similar amounts of *B. burgdorferi* cells (~ 20 μg total proteins) were subjected to SDS-PAGE, followed by immunoblots with antibodies against *B. burgdorferi* DnaK (loading control), FlaB, and FlaA. Lane 1: wild-type (WT); Lane 2: $\Delta$*flaB*$^{L}$, large loop replacement mutant; Lane 3: $\Delta$*flaB*$^{S}$, small loop replacement mutant; Lane 4: $\Delta$*flaB*, in frame deletion mutant of *flaB*; Lane 5: $\Delta$*flaB*$^{transCom}$, a trans-complemented strain of $\Delta$*flaB*.

***SI Appendix*, Table S1. Cryo-EM structure refinement statistics**

| Bacterial strains | *Borreliella burgdorferi* B31A | | *Borreliella burgdorferi* B31A | |
|---|---|---|---|---|
| Flagellar filament | Sheathed | | Non-sheathed | |
| Core flagellin | FlaB | | FlaB | |
| Sheathed protein | FlaA | | | |
| Refinement | Local | | Local | |
| PDB | 39CC | | 39CA | |
| EMDB | EMD-79303 | | EMD-79302 | |
| Resolution estimates | | | | |
| Metric | Masked (Å) | Unmasked (Å) | Masked (Å) | Unmasked (Å) |
| Using map alone (d99) | 3.90 | 3.84 | 3.62 | 3.56 |
| Overall Biso | 70.00 | 55.00 | 50.00 | 35.00 |
| d_model | 3.70 | 3.80 | 3.50 | 3.60 |
| d_model (B factors = 0) | 4.10 | 4.00 | 3.80 | 3.80 |
| FSC (model) = 0 | 3.31 | 3.38 | 3.05 | 3.15 |
| FSC (model) = 0.143 | 3.39 | 3.43 | 3.18 | 3.22 |
| FSC (model) = 0.5 | 3.87 | 7.40 | 3.71 | 4.62 |
| Mask smoothing radius (Å) | 6.82 | | 6.40 | |
| Model Validation Statistics | | | | |
| Chains | 34 | | 22 | |
| Atoms (Hydrogens) | 80464 (0) | | 49984 (0) | |
| Residues | 10522 | | 6622 | |
| RMSD bond lengths (Å) | 0.009 | | 0.014 | |
| RMSD bond angles (°) | 1.344 | | 1.732 | |
| MolProbity score | 2.16 | | 1.84 | |
| Clash score | 9.05 | | 7.82 | |
| Ramachandran favored (%) | 92.60 | | 96.24 | |
| Ramachandran allowed (%) | 6.74 | | 3.52 | |
| Ramachandran outliers (%) | 0.66 | | 0.24 | |
| Rama-Z (whole / helix / sheet / loop) | –1.26 / 0.51 / –1.08 / –2.50 | | –1.62 / –0.56 / - / –1.97 | |
| Rotamer outliers (%) | 1.90 | | 1.64 | |
| CaBLAM outliers (%) | 4.77 | | 1.71 | |
| Occupancy mean | 1.00 | | 1.00 | |
| B-factor (mean / min / max) | 107.51 / 40.82 / 312.37 | | 94.83 / 38.04 / 251.88 | |
| CC (mask / box / peaks / volume) | 0.78 / 0.49 / 0.20 / 0.78 | | 0.79 / 0.48 / 0.24 / 0.79 | |

***SI Appendix*, Table S2. Oligonucleotide primers used in this study**

| Primers | Sequences (5'-3') | Note [a] |
|---|---|---|
| P1 | ATGAGCCATATTCAACGGGA | *aph1* cassette, deletion of *flaB* [F] |
| P2 | TTAGAAAAACTCATCGAGCA | *aph1* cassette, deletion of *flaB* [R] |
| P3 | CAAGTGATATGGTTGAGGCT | 5'-flanking region of *flaB*, deletion of *flaB* [F] |
| P4 | ACGTTTCCCGTTGAATATGGCTCAT ATATCATTCCTCCATGATAAAAT | 5'-flanking region of *flaB*, deletion of *flaB* [R] |
| P5 | TTTGATGCTCGATGAGTTTTTCTAACA GATCGGCAAAAATTTAATAATTTTAG | 3'-flanking region of *flaB*, deletion of *flaB* [F] |
| P6 | GTTCTTCCTGCATTTCTCCT | 3'-flanking region of *flaB*, deletion of *flaB* [R] |
| P7 | GGATCCTGTCTGTCGCCTCTTGTGGC | 5'-flanking region of *flaB*, complementation, [F] |
| P8 | CTGCAGTTATCTAAGCAATGACAAAAC | 3'-flanking region of *flaB*, complementation, [R] |
| P9 | CTTTTTGCTCAAGAGACTGAT | Amplification of *flaA*, [F] |
| P10 | CATCAACCAAATAGCTTTCA | Amplification of *flaA*, [R] |
| P11 | AAGCTTTAGTTAAAAGCAAT | Amplification of PI-Kan, [F] |
| P12 | AAGCTTTTAGAAAAACTCAT | Amplification of PI-Kan, [R] |

a The underlined nucleotides are the engineered restriction cut sites for DNA cloning. [F] forward; [R] reverse.

**Supplementary Movie Legends**

**Movie S1. Cryo-ET reconstruction of a *B. burgdorferi* cell tip.** The tomographic reconstruction reveals periplasmic flagellar filaments with two distinct diameters.

**Movie S2. Cryo-EM structure of the half-sheathed flagellar filament.** The reconstruction reveals the FlaB filament core partially surrounded by an asymmetric FlaA sheath. Two FlaB surface loops are highlighted.

**Movie S3. Swimming behavior of Δ*flaB*$^{S}$ cells.** Real-time video microscopy reveals a nonmotile phenotype.

**Movie S4. Swimming behavior of Δ*flaB* cells.** Real-time video microscopy reveals a nonmotile phenotype.

**Movie S5. Swimming behavior of Δ*flaB*$^{L}$ cells.** Real-time video microscopy reveals reduced motility compared with wild-type cells.

**Movie S6. Swimming behavior of wild-type *B. burgdorferi* cells.** Real-time video microscopy demonstrates the characteristic motility of *B. burgdorferi*.

**Movie S7. Swimming behavior of the complemented strain.** Real-time video microscopy demonstrates restoration of motility in the complemented cells.

**Movie S8. Animation of the *B. burgdorferi* cell and its periplasmic flagella.** The animation illustrates the overall cell morphology and the organization of periplasmic flagella extending from both cell poles.
